# Environmental Control of Antimicrobial-to-Amyloidogenic Switching in Uperin 3.5 by Surfactant Assembly Dynamics

**DOI:** 10.64898/2026.03.31.715589

**Authors:** Sucharita Banerjee, Daniel Curwen, Lisandra L. Martin, Ajay Singh Panwar

## Abstract

Antimicrobial peptides (AMPs) that also form functional amyloids exhibit remarkable environmental sensitivity, yet the physicochemical rules governing their structural switching remain unresolved. Here, we investigate how surfactant charge and assembly dynamics regulate the antimicrobial–amyloidogenic transition of Uperin 3.5, a 17-residue amphibian AMP with pronounced conformational plasticity. Using an integrated approach combining all-atom molecular dynamics simulations with circular dichroism and thioflavin T fluorescence assays, we systematically probe the effects of surfactant identity, concentration relative to the critical micelle concentration (CMC), peptide stoichiometry and ionic strength. We show that α-helical stabilisation and antimicrobial-like behaviour scale directly with surfactant charge: anionic Sodium dodecyl sulphate (SDS) induces the highest helicity in monomeric Uperin 3.5 (≈80–90%), followed by zwitterionic dodecyl-phosphocholine (DPC) (≈35–45%), while cationic Cetyltrimethylammonium bromide (CTAB) fails to stabilise secondary structure. This charge-ordered trend is mirrored in oligomer remodelling, with SDS driving the most rapid dissociation of β-sheet tetramers, DPC inducing slower partial disassembly and CTAB exhibiting minimal effect. Above the CMC, micellar environments stabilise amphipathic α-helical states and efficiently dissolve amyloid assemblies. In striking contrast, under below-CMC conditions, limited SDS availability combined with peptide crowding promotes cooperative aggregation, where surfactant monomers act as dynamic scaffolds that nucleate N-terminal β-sheet interactions—an effect strongly accelerated by physiological salt. Large-scale simulations reveal mixed α/β aggregates whose formation is governed by electrostatic screening and surfactant-mediated co-assembly. Together, these findings establish surfactant charge and assembly state as quantitative, environment-dependent regulators of functional amyloidogenesis in antimicrobial peptides. More broadly, they suggest that controlled modulation of membrane-mimetic environments can be exploited to bias peptides toward antimicrobial or amyloidogenic states, offering conceptual avenues for therapeutic strategies targeting peptide misfolding and neurodegenerative disorders.

## Introduction

Antimicrobial peptides (AMPs) are an evolutionarily conserved component of innate immunity, exhibiting broad-spectrum activity against a range of pathogens^1^. Beyond their canonical antimicrobial function, a subset of AMPs—including Uperin 3.5—have been found to possess amyloidogenic properties, enabling them to adopt β-sheet-rich aggregates under specific environmental conditions^2,3^. This dual functionality has sparked growing interest in understanding the environmental factors that regulate their conformational states and biological roles^4,5^.

Uperin 3.5, a 17-residue peptide originally isolated from the skin of the Australian toadlet *Uperoleia mjobergii*, exemplifies this structural plasticity^6–8^. Depending on its microenvironment, the peptide can exist in a disordered, α-helical, or β-sheet-rich amyloid state^9–11^. This polymorphism is hypothesised to underlie its ability to toggle between antimicrobial and amyloid or structural roles, a phenomenon now recognized as functional amyloidogenesis^5^. Unravelling the factors that influence these conformational transitions is therefore critical to both understanding its biological function and harnessing its potential in biomedical applications^12^.

One such factor is the presence of membrane-mimetic environments, such as surfactants, which emulate the physicochemical properties of lipid bilayers^13,14^. Surfactants, such as sodium dodecyl sulphate (SDS), dodecyl phosphocholine (DPC), and cetyltrimethylammonium bromide (CTAB) differ in their head group charges—anionic, zwitterionic, and cationic, respectively—providing distinct electrostatic and hydrophobic environments that can influence peptide conformation, oligomerisation and aggregation^15,16^. Additionally, surfactant concentration relative to the critical micelle concentration (CMC) further modulates the local microenvironment, with monomeric and micellar forms offering different interaction interfaces to peptides^17^.

In this study, we employ an integrated approach combining all-atom Molecular Dynamics (MD) simulations and experimental validation to systematically investigate how surfactant identity (SDS, DPC, CTAB) and concentration (below and above CMC) influence the conformational behaviour and aggregation propensity of Uperin 3.5. In addition, for SDS below-CMC systems, we examined the effect of salt on aggregation by comparing peptide ensembles simulated with and without added ions^18^. We simulate both monomeric and tetrameric forms of the peptide under these conditions, allowing us to dissect the early events in aggregation and secondary structure evolution^19^. Complementary circular dichroism (CD) spectroscopy^20^ and thioflavin-T (ThT) fluorescence assays^21^ provide experimental confirmation of surfactant-induced folding and aggregation transitions observed *in silico*.

Our results reveal that surfactant type, concentration and ionic conditions distinctly modulate the conformational ensemble and self-assembly of Uperin 3.5, offering mechanistic insight into its functional versatility and highlighting the importance of membrane mimetics in amyloidogenic peptide research.

## Materials and methods

### Experimental methods

#### Materials

Uperin 3.5 peptide with a C-terminal amide (>98% purity) was purchased from Peptide 2.0 Inc. Sodium dodecyl sulfate (SDS) and dodecylphosphocholine (DPC) were obtained from Pall Life Sciences. Thioflavin T (ThT) and buffer components were purchased from Sigma-Aldrich. All solutions were prepared using ultrapure water (18.2 MΩ·cm). Phosphate-buffered saline (PBS) contained 20 mM phosphate and 100 mM NaCl at pH 7.4.

#### Circular Dichroism Spectroscopy

Circular dichroism (CD) measurements were performed using a Jasco J-815 spectropolarimeter equipped with a temperature-controlled cuvette holder set to 37 °C. Peptide samples were prepared at a final concentration of 100 μM following addition of PBS and surfactant solutions. Spectra were collected between 190–260 nm (or ≥195 nm in PBS) using a 1 mm pathlength quartz cuvette. Time-course measurements were recorded over 24 h to monitor secondary structure evolution. Raw spectra were baseline corrected and converted to mean residue molar ellipticity for analysis.

#### ThT Fluorescence Assays

Peptide aggregation was monitored using ThT fluorescence in black 96-well microplates using a CLARIOstar plate reader (BMG Labtech). Measurements were performed at 37 °C with excitation and emission wavelengths of 440 nm and 480 nm, respectively, and fluorescence intensities were recorded at 10 min intervals. Two experimental protocols were employed. In the first, surfactants were introduced prior to PBS addition to examine peptide aggregation in the presence of surfactants. In the second, peptides were first allowed to aggregate in PBS, after which surfactants were added to probe their effect on pre-formed aggregates.

#### Molecular Dynamics Simulations

All-atom molecular dynamics simulations were performed using NAMD^22^ with the CHARMM36m^23^ force field for proteins and lipids and the TIP3P water model^24^. The setups were simulated at 310 K and 1 atm under periodic boundary conditions with long-range electrostatics treated using the particle mesh Ewald method^25,26^. The simulations were performed to investigate the interactions between Uperin 3.5 (U3.5) peptides—both as monomers and tetramers—and three surfactants of differing charge characteristics: anionic sodium dodecyl sulphate (SDS), zwitterionic dodecyl phosphocholine (DPC), and cationic cetyltrimethylammonium bromide (CTAB)^27–30^. Two classes of peptide–surfactant conditions were investigated. In the below-CMC simulations, monomeric or tetrameric Uperin 3.5 peptides were surrounded by dispersed surfactant monomers (12 SDS, 8 DPC, or 8 CTAB molecules) randomly distributed around the peptides. In micelle-based simulations, peptides were positioned near the surface of preformed surfactant micelles composed of 60 SDS^31,32^ or DPC molecules^33^ or 120 CTAB molecules^34^, corresponding to experimentally reported micelle sizes. These systems enabled the investigation of peptide interactions with both dispersed surfactants and micellar assemblies. To examine aggregation behaviour under crowded conditions, additional simulations containing 20 peptides were constructed. Two ensembles were considered: a non-aggregated system comprising 20 peptides initially in random-coil conformations in the presence of SDS monomers under salt-free conditions, and a partially aggregated system consisting of two preformed tetramers together with 12 monomeric peptides in the presence of SDS and 0.15 M NaCl. These simulations were used to assess the influence of ionic strength on peptide aggregation under below-CMC surfactant conditions. Further simulation details and setup parameters are provided in the Supplementary Information (Tables ST1 and ST2).

## Results and Discussion

### Dynamic surfactant co-assembly under below-CMC conditions drives rapid, charge-dependent peptide remodelling, while micelles stabilise α-helices with slower kinetics

To determine how surfactant charge and assembly state influence the conformational behaviour of Uperin 3.5, all-atom molecular dynamics simulations were performed for both monomeric and tetrameric peptide forms in the presence of three surfactants: anionic SDS, zwitterionic DPC, and cationic CTAB. The simulation setups were examined under conditions below and above the respective critical micelle concentrations (CMCs), representing environments dominated by dispersed surfactant monomers and pre-formed micelles, respectively. The initial simulation configurations for each case are shown in **Figure 1**, while **Figure 2** presents representative final peptide–surfactant assemblies together with the time-resolved evolution of peptide secondary structure.

**Figure 1.**
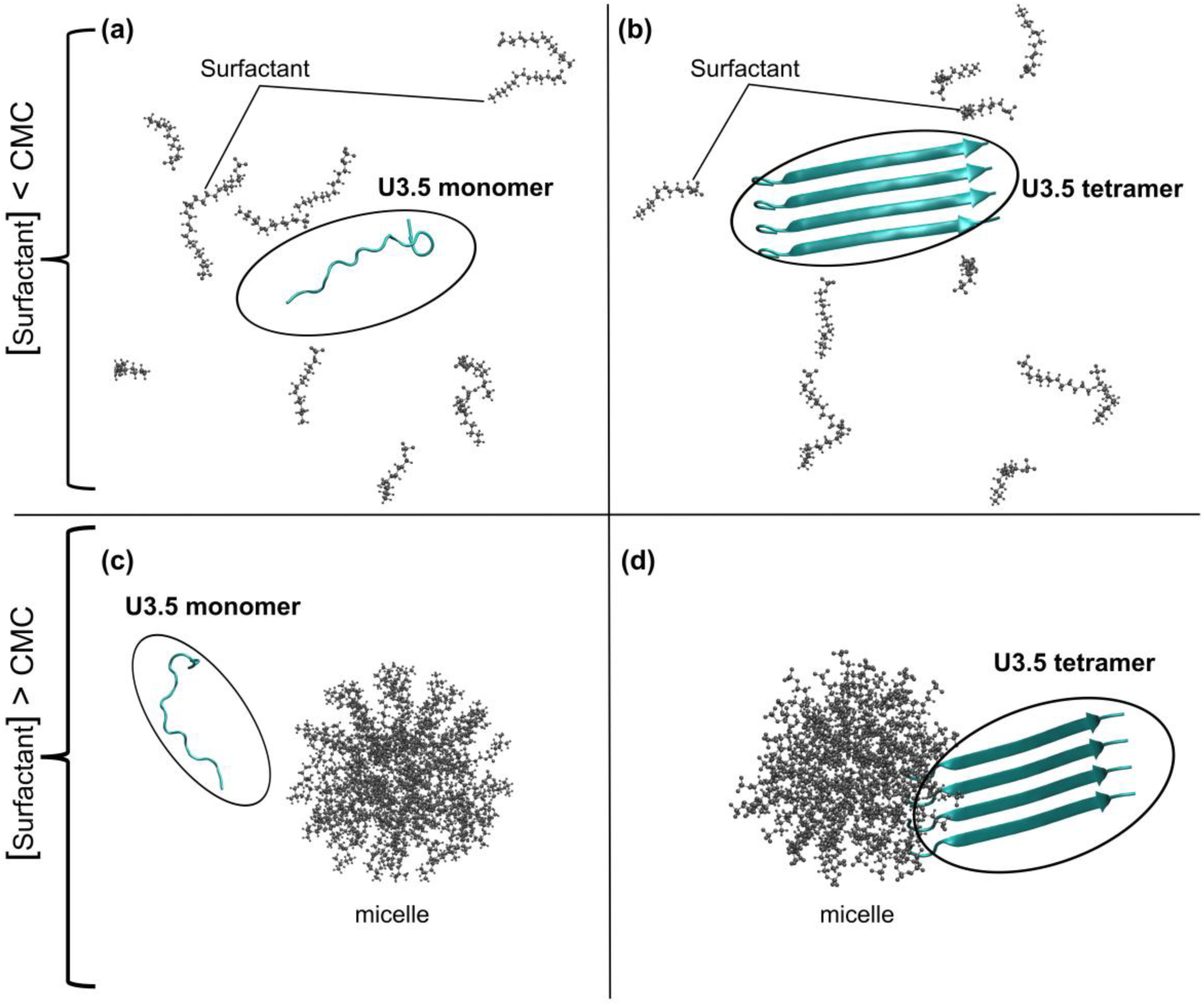
Schematic representation of Uperin 3.5 peptide–surfactant simulation setups; snapshots correspond to the initial configuration (t = 0). (a)Monomeric peptide surrounded by dispersed surfactant molecules (SDS, DPC, or CTAB) under sub-critical micelle concentration (below-CMC) conditions.(b)Tetrameric peptide assembly interacting with dispersed surfactant molecules under below-CMC conditions. (c)peptide monomer positioned near a preformed surfactant micelle, representing above-CMC conditions.(d) Tetrameric peptide assembly interacting with a surfactant micelle under above-CMC conditions.

**Figure 2.**
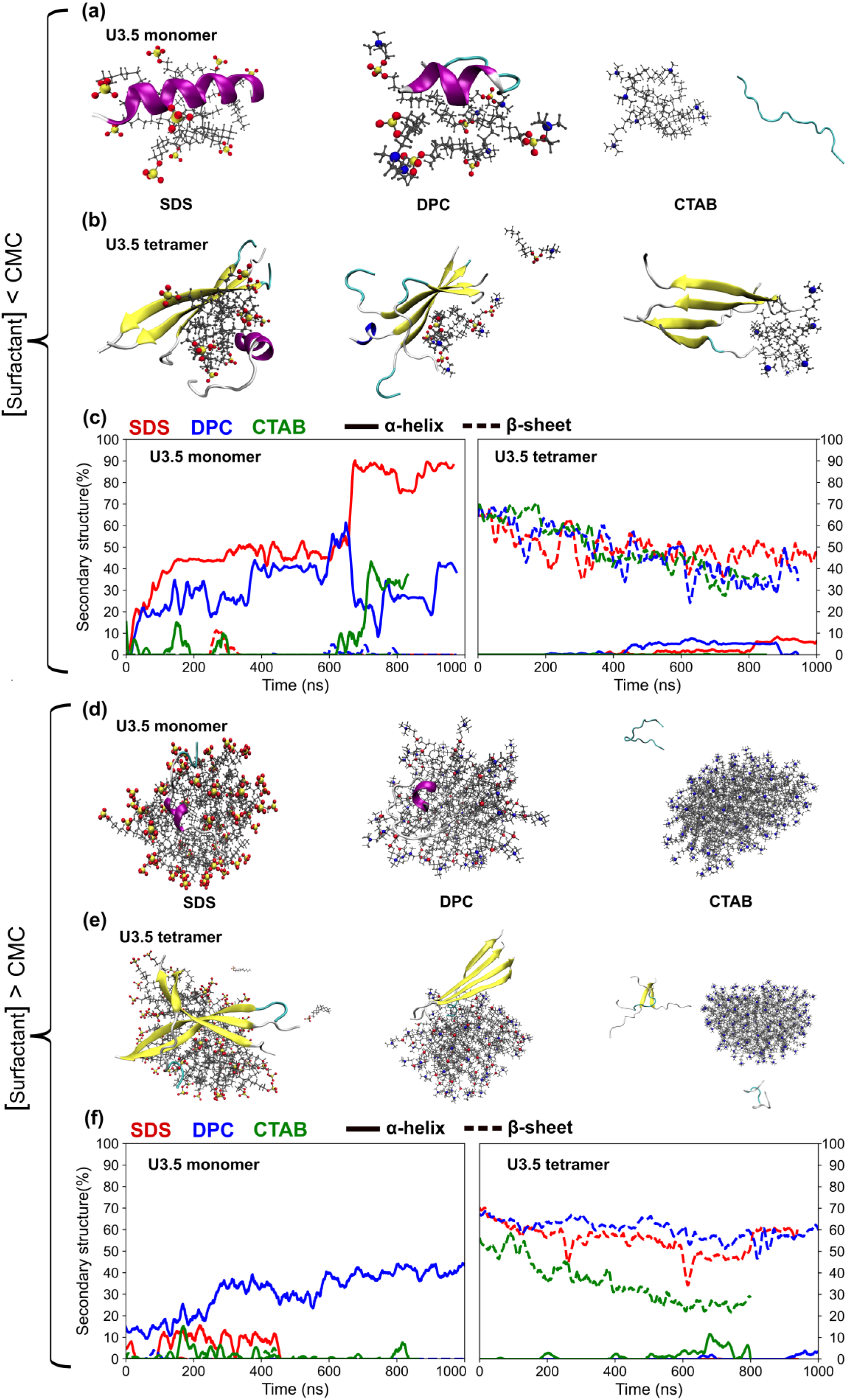
Molecular dynamics simulations of Uperin 3.5 peptides interacting with surfactants under below-CMC and micelle-based conditions.(a,b) Final simulation snapshots of monomeric (a) and tetrameric (b) Uperin 3.5 peptides interacting with dispersed surfactant monomers (SDS, DPC, or CTAB) under below-CMC conditions.(c) Time evolution of secondary structure content for monomeric and tetrameric Uperin 3.5 peptides in the presence of surfactant monomers.(d,e) Final simulation snapshots of monomeric (d) and tetrameric (e) Uperin 3.5 peptides interacting with preformed surfactant micelles.(f) Time evolution of secondary structure content for monomeric and tetrameric Uperin 3.5 peptides in micelle-based setups.

Under below-CMC conditions (**Figure 1a,b**), surfactant molecules were initially (t_0_) distributed as individual monomers around the peptide. During the simulations, these monomers dynamically reorganised around the peptide, forming transient pseudo-micellar assemblies that strongly influenced peptide structure. In the presence of SDS, the monomeric peptide rapidly adopted a highly stable α-helical conformation (**Figure 2a**), with helicity exceeding ∼90% over the trajectory (**Figure 2c**). This behaviour is consistent with favourable electrostatic interactions between the negatively charged sulfate headgroups of SDS and the cationic residues of Uperin 3.5, which anchor the peptide at the surfactant interface while hydrophobic tail interactions stabilise the amphipathic helix.

In contrast, DPC monomers formed a less compact and more weakly organised environment around the peptide (**Figure 2a**). Consequently, only partial helix induction was observed, with α-helical content stabilising near ∼40% over the simulation (**Figure 2c**). The reduced efficiency relative to SDS likely reflects weaker electrostatic anchoring arising from the overall neutral charge of DPC, leading to less persistent peptide–surfactant interactions. Cationic CTAB monomers failed to establish stable associations with the peptide (**Figure 2a**), and little or no secondary structure development was observed throughout the trajectory (**Figure 2c**), consistent with electrostatic repulsion between the positively charged headgroups and the cationic peptide.

The formation of α-helical structure in Uperin 3.5 is strongly influenced by the surrounding environment. In aqueous solution, peptide–water interactions compete with intramolecular backbone hydrogen bonding and can destabilize ordered secondary structures. Upon interaction with surfactant assemblies, hydrophobic residues preferentially partition toward the micellar interior while polar residues remain solvent exposed, reducing backbone solvation and promoting stabilizing intramolecular *i → i+4* hydrogen bonds. Similar environmental effects have been reported in membrane-mimetic solvents such as trifluoroethanol (TFE), where reduced peptide solvation promotes helix formation and peptide self-assembly^35^. These observations highlight the role of amphiphilic environments in stabilizing α-helical conformations in antimicrobial peptides.

The tetrameric peptide displayed related but structurally distinct behaviour under below-CMC conditions (**Figure 1b**). In SDS-containing s, the initially β-sheet-rich tetramer underwent progressive destabilisation of inter-peptide hydrogen bonding, accompanied by partial dissociation and the emergence of α-helical segments (**Figure 2b,c**). This behaviour indicates that dynamically assembling SDS monomers are capable not only of stabilising helical conformations but also of actively disrupting amyloidogenic β-sheet contacts. DPC induced similar but less extensive remodelling, whereas CTAB again produced negligible structural changes, reinforcing the importance of electrostatic complementarity in enabling peptide restructuring in the monomeric surfactant regime.

A distinct behaviour emerged above the CMC, where surfactants were present as pre-assembled micelles (**Figure 1c,d**). In SDS and DPC cases, the monomeric peptide rapidly adsorbed onto the micelle surface and adopted stable α-helical conformations (**Figure 2d**), as reflected by sustained helicity over the simulation trajectory (**Figure 2f**). The amphipathic peptide was oriented such that hydrophobic residues interacted with the micelle core while polar and charged residues remained solvent exposed, consistent with membrane-mimetic binding. However, despite the higher surfactant concentration, the kinetics of helix formation were slower than those observed under below-CMC conditions.

Tetrameric peptides interacting with micelles exhibited partial destabilisation of the β-sheet assembly but remained largely associated over the simulation timescale (**Figure 2e,f**). In both SDS and DPC setups, interactions with the micelle surface induced modest structural rearrangements and limited α-helix formation, yet complete tetramer dissociation was not observed. As in the monomer simulations, CTAB micelles failed to support productive peptide association or structural transitions.

The slower remodelling kinetics observed in micellar conditions can be rationalised by fundamental differences in surfactant assembly dynamics. Below the CMC, individual surfactant monomers remain highly mobile and can dynamically reorganise around the peptide, enabling cooperative co-assembly, efficient hydrophobic shielding, and rapid disruption of inter-peptide β-sheet contacts. In contrast, above the CMC surfactants are sequestered within stable micelles, and peptide interactions are largely restricted to surface adsorption. The structural rigidity and curvature constraints of micelles limit surfactant rearrangement and peptide penetration into the hydrophobic core, thereby imposing kinetic barriers to extensive conformational remodelling.

Collectively, these results demonstrate that peptide structural fate is governed not simply by surfactant concentration but by the dynamic assembly state and charge of the surfactant environment. Below the CMC, flexible co-assembly with individual surfactant monomers enables rapid, charge-dependent peptide remodelling, promoting helix formation and destabilising β-sheet aggregates. In contrast, micellar environments above the CMC stabilise α-helical conformations but impose kinetic constraints on peptide restructuring. This distinction highlights a mechanistic continuum in which dynamic peptide–surfactant co-assembly promotes rapid structural switching, whereas pre-formed micelles favour stable but less adaptable antimicrobial-like states.

### Energetic determinants of peptide–surfactant recognition reveal electrostatic dominance

To gain mechanistic insight into how different surfactants modulate the structure and stability of Uperin 3.5, quantitative analyses of intermolecular interactions were performed using radial distribution functions (RDFs), hydrogen bonding patterns, and radius of gyration (*Rg*) measurements. These metrics revealed how electrostatics, hydrophobicity, and structural flexibility influence peptide folding and aggregation in the presence of SDS, DPC and CTAB.

RDFs were calculated to compare the spatial distribution of surfactant head groups and hydrophobic tails relative to key residues of the peptide (**Figure 3a, b**). The results revealed a consistent trend across both monomeric and tetrameric peptide simulations: SDS exhibited the strongest and most localised interaction with the peptide, followed by DPC, with CTAB showing minimal interaction (**Figure 3a**). This pattern is attributed to the electrostatic complementarity between the negatively charged sulphate headgroups of SDS and the positively charged residues of Uperin 3.5, which facilitate strong peptide–surfactant association. In contrast, DPC, being zwitterionic, lacks a net charge, resulting in moderate interaction strength. CTAB, carrying a positively charged quaternary ammonium group, repels the cationic residues of the peptide, preventing stable complex formation. Similar trends were observed in the RDFs describing interactions between the hydrophobic tails of the surfactants and hydrophobic regions of the peptide (**Figure 3b**). SDS again showed the highest interaction probability, indicating deeper insertion of peptide hydrophobic residues into the surfactant tail region, consistent with helix formation. DPC showed moderate tail-residue association, while CTAB’s interactions were minimal due to the lack of peptide association in the first place. These RDF profiles align well with previously observed secondary structure trends, and underscore the importance of both electrostatic and hydrophobic interactions in driving peptide folding and aggregation.

**Figure 3.**
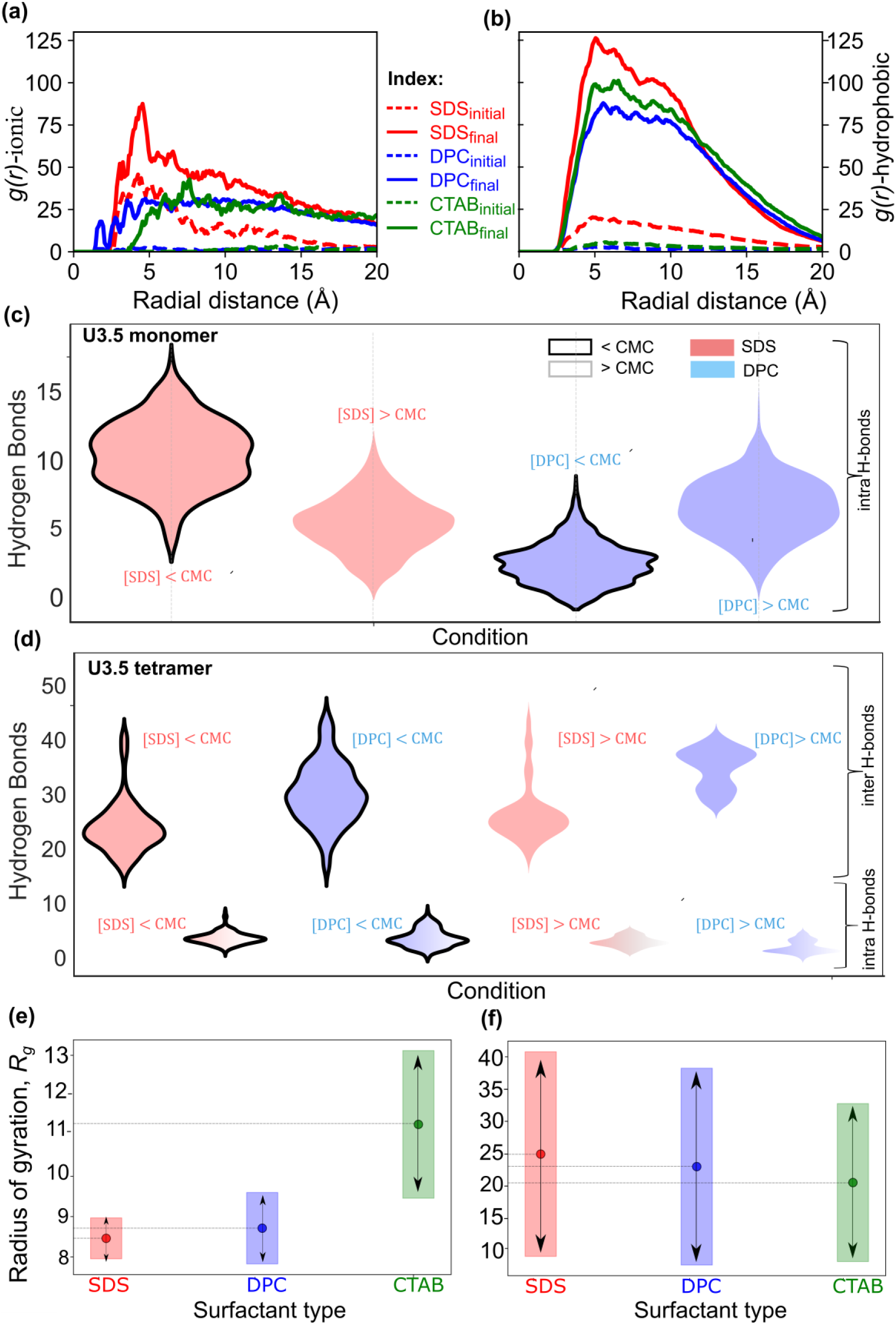
(a) Radial Distribution Function (RDF, *Rg*) analysis depicting the interactions between the charged head groups of surfactants and the corresponding oppositely charged residues of the peptides. (b) RDF analysis of hydrophobic interactions between the surfactant tail groups and the hydrophobic residues of the peptides. (c) Number of intramolecular hydrogen bonds formed by the peptide monomer in the presence of SDS and DPC, at surfactant concentrations below and above the critical micelle concentration (CMC). (d) Variation in intra- and intermolecular hydrogen bond formation within the peptide tetramer upon interaction with SDS and DPC under below-CMC and above-CMC conditions. (e, f) Changes in the radius of gyration of the peptide monomer (e) and tetramer (f) upon interaction with the three surfactants of differing charge characteristics.

Intra-peptide hydrogen bonding within the peptide was analysed as a marker of structural integrity and secondary structure formation. In the monomer simulations (**Figure 3c**), SDS promoted the highest number of intramolecular hydrogen bonds, consistent with the formation of stable α-helical structures. DPC also supported intra-peptide hydrogen-bonding (H-bonding), albeit to a lesser extent, while CTAB systems exhibited minimal hydrogen bonding, reflecting the lack of secondary structure development. Interestingly, in below-CMC conditions, the flexible and adaptive nature of freely dispersed surfactant monomers enabled faster peptide wrapping and H-bond stabilisation compared to micelle systems. This supports the hypothesis that below-CMC conditions can lead to more dynamic and cooperative folding environments due to the greater conformational freedom of individual surfactant molecules. In the tetramer simulations (**Figure 3d**), hydrogen bond analysis provided a quantitative view of β-sheet disassembly. The number of inter-peptide hydrogen bonds—a proxy for β-sheet integrity—decreased most rapidly in the presence of SDS, followed by DPC. This reflects the ability of these surfactants to destabilise amyloidogenic β-sheet assemblies. At the same time, a gradual increase in intramolecular hydrogen bonds was observed, indicating the onset of helix formation as β-sheets dissolved. These changes were slower in DPC, while CTAB systems retained a high inter-peptide H-bond count, indicating that the tetramer remained structurally stable throughout. Taken together, these results reinforce a charge- and concentration-dependent mechanism in which surfactants disrupt amyloid structures and promote α-helical conformations through distinct interaction pathways. The detailed hydrogen-bond analysis (**Figure S1**) over the trajectories supports these trends, demonstrating that SDS and DPC modulate intra- and intermolecular hydrogen bonding in a concentration-dependent manner.

To assess changes in overall peptide compactness, we monitored the radius of gyration (*Rg*) of both monomeric (**Figure 3e**) and tetrameric (**Figure 3f**) cases. For the monomer, SDS induced the most compact conformation (lowest *Rg*), consistent with the formation of a tightly folded helix stabilized by both hydrophobic collapse and electrostatic interactions. DPC showed intermediate compaction, while CTAB exhibited the highest *Rg* values, reflecting the lack of structural consolidation. For the tetramer, a contrasting trend was observed. The presence of SDS led to increased fluctuations in *Rg*, indicative of structural rearrangement and partial disassembly of the β-sheet core. DPC caused moderate destabilization, while CTAB had minimal impact, with *Rg* remaining relatively stable over time. These observations correlate well with the hydrogen bonding analysis, where SDS and DPC disrupted inter-strand interactions and induced local unfolding, while CTAB preserved the native β-sheet conformation. Consistent with this interpretation, the detailed *Rg* analysis over time (**Figure S2**) showed that SDS stabilizes the monomer most strongly while disrupting tetramer compactness, whereas DPC exerts intermediate effects and CTAB shows minimal interaction. Overall, the *Rg* data corroborate our mechanistic model: surfactants with higher charge complementarity and stronger hydrophobic interactions (SDS > DPC > CTAB) not only influence peptide secondary structure but also modulate overall structural dynamics and aggregation state.

While the above analyses focused on isolated monomeric and tetrameric systems to elucidate the fundamental modes of peptide–surfactant interaction, real biological and experimental environments often involve much higher local peptide concentrations, where multiple peptides can simultaneously interact with a limited number of surfactant molecules. Indeed, our ThT fluorescence experiments indicated that under below-CMC SDS conditions, particularly when peptide concentrations were high and surfactant availability was limited, significant aggregation was observed—contrasting the helix-promoting behaviour seen at low peptide densities. These observations raised important questions about how peptide crowding and surfactant scarcity might cooperatively modulate aggregation behaviour. To investigate these effects in greater mechanistic detail, we performed additional molecular dynamics simulations involving ensembles of 20 peptides in the presence of below-CMC SDS concentrations, with and without added salt. These simulations allowed us to mimic the experimental conditions more closely and to dissect the underlying molecular mechanisms driving aggregation under physiologically relevant crowded conditions.

### Peptide crowding and limited surfactant availability below the CMC drive amyloidogenic aggregation via SDS-mediated clustering

To explore how surfactant concentration and peptide crowding influence aggregation behaviour, we conducted both experimental (Thioflavin T, ThT assays) and computational (molecular dynamics) investigations under below-CMC SDS conditions. These studies revealed that peptide aggregation is not solely dependent on surfactant identity and charge but is strongly modulated by the relative stoichiometry between peptides and surfactants, as well as by the ionic strength of the medium.

Two experimental protocols^10,35–37^ were employed to evaluate peptide aggregation in the presence of SDS and DPC. In the first protocol, peptides were introduced into SDS- or DPC-containing solutions (below- or above-CMC) without any added salt. These samples generally showed no measurable ThT fluorescence, indicating an absence of β-sheet-rich aggregates. This suggests that the high surfactant-to-peptide ratio favoured helix induction and suppressed amyloid formation. In the second protocol, peptides were first incubated in PBS buffer (with physiological salt), allowing partial or complete aggregation before the addition of surfactants. Interestingly, even after pre-formed aggregation, the subsequent introduction of SDS or DPC led to disassembly of aggregates and a transition toward α-helical states, as indicated by a reduction in ThT signal over time. This behaviour supports a functional switch toward antimicrobial conformations, as previously observed in membrane-mimetic systems.

However, a distinct deviation was observed for SDS under below-CMC conditions, where both experimental protocols showed increased ThT fluorescence, indicative of enhanced aggregation or β-sheet clustering. This suggests that, under specific concentration and ionic conditions, SDS monomers may paradoxically promote aggregation, contrary to the expected disassembly behaviour observed at higher concentrations.

To validate the molecular mechanism underlying these observations, we designed two MD simulation setups mimicking the experimental ThT protocols under below-CMC SDS conditions. In the non-aggregated condition, 20 Uperin 3.5 peptides—each initially in a random coil conformation—were simulated in pure water containing 12 SDS monomers (i.e., below CMC), and no added salt. Over the course of 1 μs, we observed the formation of multiple small peptide clusters, each comprising 6–8 peptides. Interestingly, these clusters were heterogeneous in nature—some composed of β-sheet structures, others of α-helices, and some comprising a mix of both. The SDS monomers appeared to serve as nucleation centres or scaffolds, clustering locally and screening peptide charges, thus enabling aggregation. This dynamic and non-specific clustering behaviour was not observed in monomeric simulations, underscoring the influence of peptide crowding and limited surfactant availability. In the aggregated condition, designed to mimic the second ThT protocol, we simulated 20 peptides, including two pre-formed β-sheet tetramers and 16 random coil monomers in the presence of 12 SDS monomers and 0.15 M NaCl. Here, we observed rapid and extensive aggregation, with nearly all peptides forming a single large cluster within the first few hundred nanoseconds. The final aggregate exhibited mixed secondary structure, consisting of both β-sheets and helices stabilised around a central SDS cluster. This behaviour was notably more cooperative and faster than in the salt-free ensemble, highlighting the role of ionic screening in promoting inter-peptide associations.

Snapshots of the initial and final configurations for both systems are shown in **Figure 4a, b**, alongside their corresponding ThT assay results. **Figure 5a** provides a quantitative analysis of peptide clustering over time, clearly demonstrating a single large aggregate in the salt-containing model versus three smaller clusters in the salt-free model.

**Figure 4.**
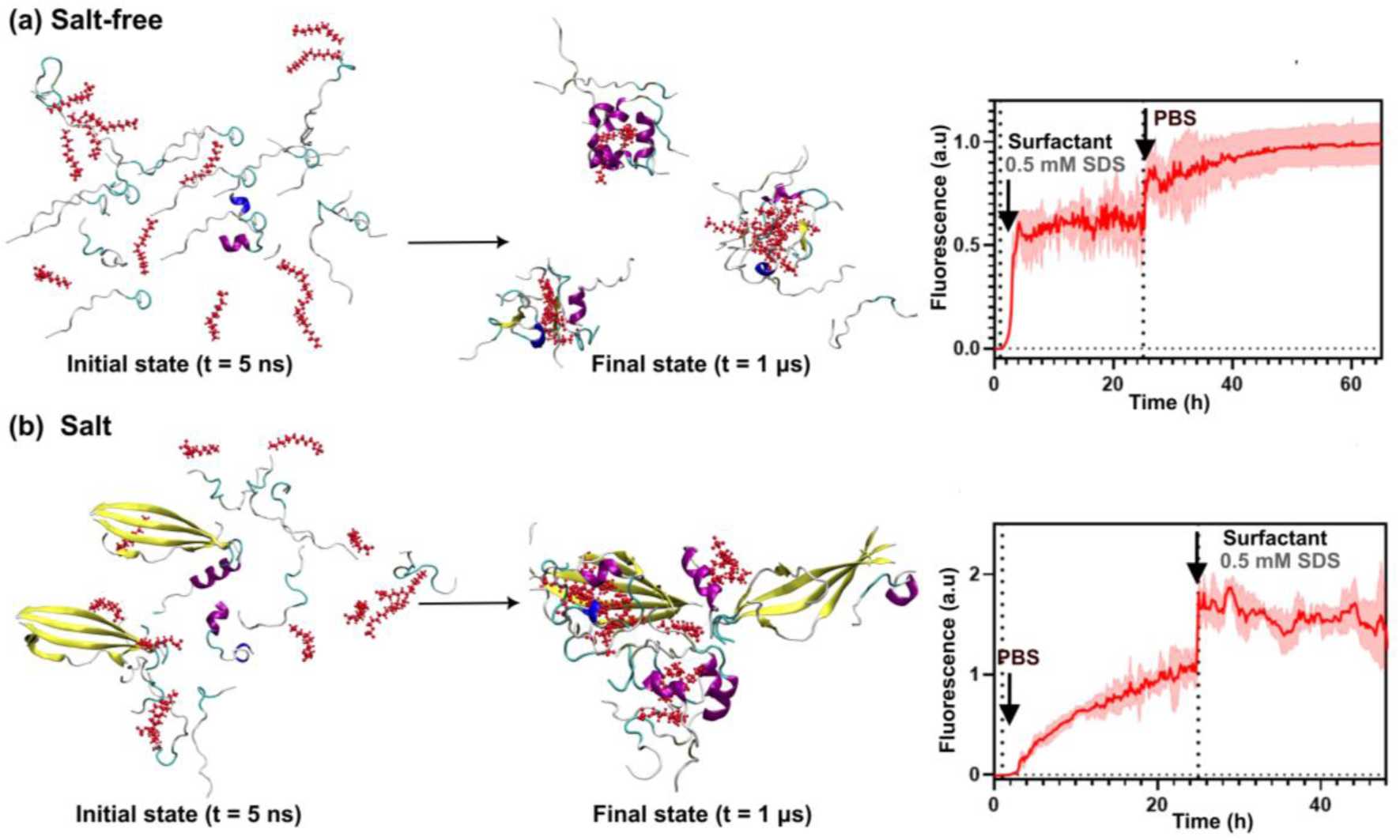
(a) Initial and final simulation snapshots of a salt-free system containing 20 non-aggregated, random coil Uperin 3.5 peptides in the presence of SDS monomers at below-CMC concentrations.Corresponding Thioflavin T (ThT) fluorescence assay results from an experimental condition where SDS (below CMC) was added directly to peptides in aqueous solution, followed by the addition of phosphate-buffered saline (PBS)(b) Initial and final simulation snapshots of a system containing 20 peptides—comprising a mixture of pre-formed β-sheet tetramers and random coil peptides—simulated under similar SDS conditions but in the presence of salt. Corresponding ThT assay results reflect the experimental setup where peptide aggregation was first induced by PBS addition, followed by the introduction of SDS monomers at below-CMC concentrations.

**Figure 5.**
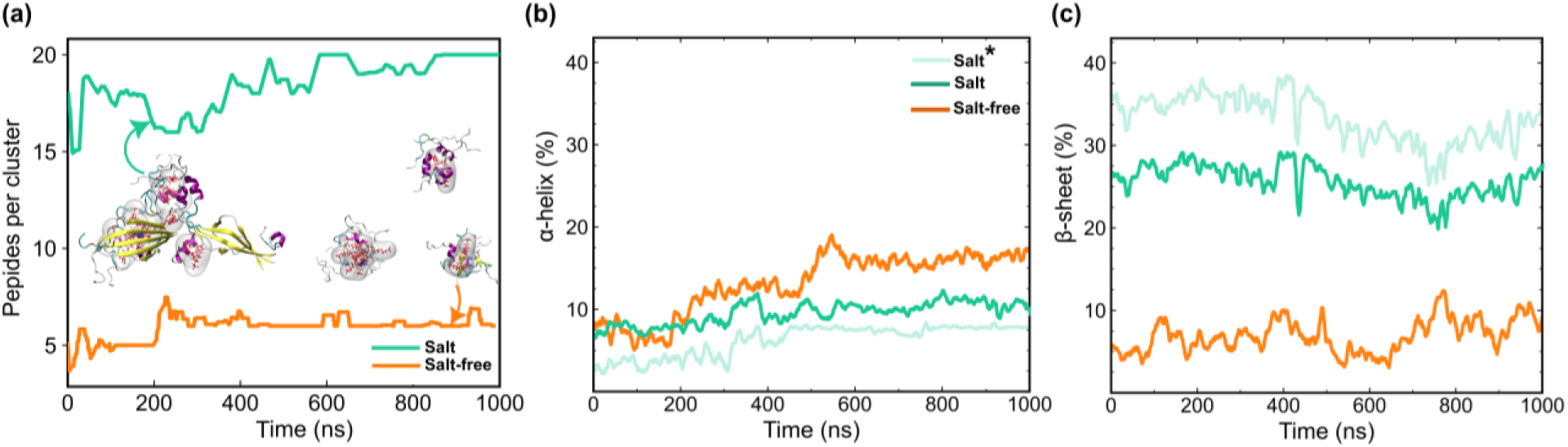
Comparative analysis of aggregation and secondary structure evolution in 20 Uperin 3.5 peptide–SDS below CMC ensembles with and without salt. (a) Aggregate size distribution as a function of simulation time, showing the number of peptides per aggregate, where an aggregate is defined by a 5 Å inter-peptide distance cutoff. The 20-peptide ensemble in the presence of 0.15 M NaCl exhibits the formation of a single large aggregate, whereas the salt-free condition shows multiple smaller aggregates. The average evolution of (b) α-helical and (c) β-sheet content is shown across the simulation trajectories. In each panel, total secondary structure content is shown for both salt and salt-free conditions, alongside a separate trace (Salt*, light sea green) representing the salt condition excluding contributions from the initially present β-sheet tetramers. This distinction highlights the *de novo* formation of secondary structure in the remaining peptides over time, enabling direct assessment of structural induction rather than propagation from preformed aggregates.

Secondary structure evolution for individual peptides, shown as heat maps in **Figure S3**, revealed that the salt-free condition exhibited a modest increase in α-helical content, particularly within the smaller clusters, suggesting a balance between aggregation and folding. In contrast, the salt-containing ensemble displayed a complex behaviour: retention of β-sheet content from the initial tetramers combined with new helix formation, resulting in a hybrid aggregate. The observed aggregation under below-CMC SDS conditions may be explained by several factors. At low surfactant concentrations, SDS monomers are insufficient to fully coat or solubilize all peptides, leading to partial surfactant-mediated screening of charges. This can enable short-range interactions and clustering of peptides, especially under crowded conditions. SDS monomers act as focal points for aggregation, enhancing local peptide concentration and facilitating inter-peptide β-sheet or helical stacking, depending on residue exposure and orientation. The presence of salt enhances electrostatic screening, reducing repulsion among positively charged residues and accelerating aggregate formation. A further detailed time-resolved secondary-structure profiles for all peptides in the 20-peptide setups that confirm the divergent aggregation pathways under salt and salt-free conditions has been depicted in **Figure S4**.

Together, these results demonstrate that below-CMC SDS concentrations can paradoxically enhance aggregation in crowded peptide systems, especially when the peptide-to-surfactant ratio is high, and salt is present. This illustrates the strong dependence of peptide behaviour on environmental context: under micellar (above-CMC) conditions, SDS stabilises helical conformations that support antimicrobial activity, whereas under below-CMC conditions with high peptide density and salt, the same surfactant instead promotes β-sheet aggregation. Thus, the structural and functional outcome is not an intrinsic property of the peptide alone but is dictated by the specific balance of surfactant concentration, peptide stoichiometry, and ionic strength.

While the overall aggregation behaviour and secondary structure evolution provided important insights into the macroscale outcomes of peptide crowding under below-CMC SDS conditions, we next sought to characterise the molecular details of these aggregation processes. Specifically, we aimed to identify which regions of the peptides mediate inter-peptide interactions during cluster formation, and how these interactions evolve over time. To this end, we performed residue–residue contact map analyses on the 20-peptide simulation trajectories. Additionally, we examined electrostatic and van der Waals energy contributions to further quantify the thermodynamic stability of the aggregates. Together, these analyses provide a more granular understanding of the driving forces underlying SDS-facilitated peptide clustering.

### Ionic strength accelerates β-sheet nucleation and drives cooperative aggregate formation

To further elucidate the nature and progression of peptide–peptide interactions in crowded environments under below-CMC SDS conditions, residue–residue contact map analyses were performed on both salt-free and salt-containing 20-peptide systems. These maps offer a detailed visualisation of how specific regions of Uperin 3.5 engage in inter-peptide contacts over the course of the simulation, complementing structural and clustering analyses.

### N-terminal interaction hotspots mediate early-stage aggregation across conditions

In the aggregated system containing salt (**Figure 6a**), contact maps of the early-stage (initial) and late-stage (final) configurations revealed key insights into the aggregation mechanism. At the initial time point, prominent contacts were already observed among the N-terminal regions of the peptides, suggesting preferential interactions in this segment even at early stages of assembly. As the simulation progressed, contact density increased and extended across the entire peptide length, indicating the formation of tighter, more stabilised inter-peptide networks. This pattern is consistent with a scenario in which initial nucleation occurs via the N-terminal regions, followed by structural consolidation and maturation into a large, mixed-content aggregate.

**Figure 6.**
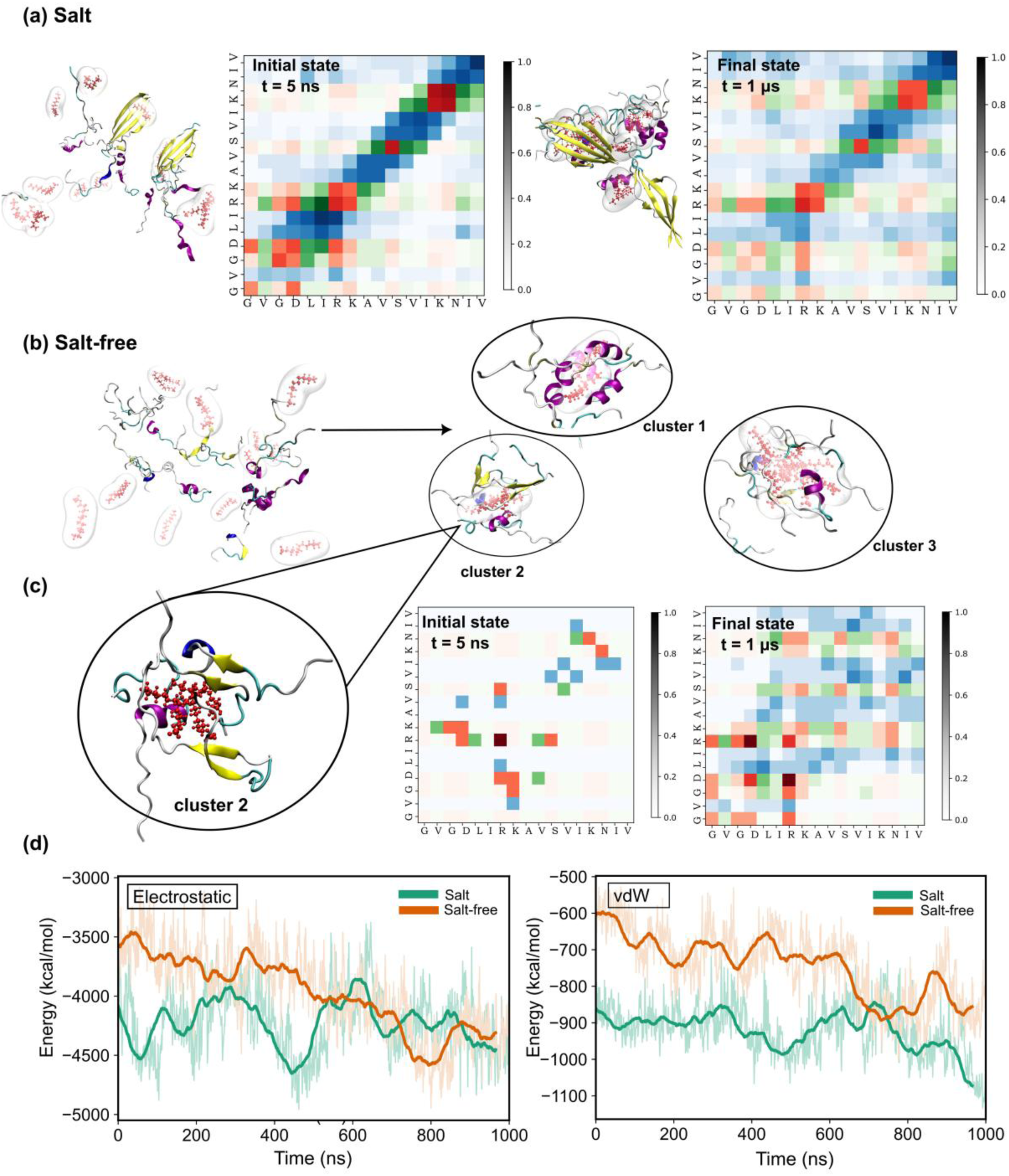
Inter-peptide contact maps and corresponding initial and final snapshots of aggregated and non-aggregated peptide systems simulated in salt-containing and salt-free environments, respectively. Secondary structure evolution was analysed to investigate residue–residue interactions associated with structural transitions. (a) Initial and final snapshots of the system simulated in the presence of salt, along with corresponding residue–residue contact maps, highlighting the progression of aggregation. (b) In the salt-free system, partial aggregation into small clusters was observed at the final stage. (c) Cluster 2 from this system is highlighted along with its contact map, which reveals interaction patterns similar to those observed in the salt-containing system but with comparatively slower aggregation kinetics. In both systems, strong interactions were predominantly localised near the N-terminus, likely corresponding to the formation of β-sheet-rich aggregates. Hydrophilic and hydrophobic residues are depicted in red and blue, respectively. Residue–residue contact maps a–c follow a colour-coded scheme: red (hydrophilic–hydrophilic), blue (hydrophobic–hydrophobic), and green (hydrophilic–hydrophobic), with colour intensity indicating the frequency of contacts. (d) Comparison of electrostatic and van der Waals energy profiles between the two conditions demonstrates that both conditions promote aggregation, with the salt-containing ensemble exhibiting faster kinetics. In both cases, the presence of SDS monomers below the critical micelle concentration (CMC) facilitates aggregation, ultimately driving the systems toward energetically favourable states.

For the salt-free system (**Figure 6b**), a representative cluster (cluster two) peptide clusters, each comprising 6-7 peptides, formed along the trajectory, each comprising either helices, beta sheets or a hybrid of both, out of which one has been selected) was analysed (**Figure 6c**). This cluster, which adopted predominantly β-sheet-rich features at the end of the trajectory, also showed limited initial contacts but developed strong interactions among the N-terminal residues over time. Despite being slower and less extensive, the contact distribution pattern mirrored that of the salt-containing system, reinforcing the role of N-terminal residues as aggregation hotspots, likely due to their high β-sheet propensity and involvement in early nucleation events.

This recurring N-terminal interaction signature in both systems suggests that SDS-mediated aggregation under below-CMC conditions follows a conserved structural pathway, driven initially by directional hydrogen bonding and strand pairing among aggregation-prone residues in the peptide’s N-terminus. The detailed inter-peptide contact maps for both salt and salt-free systems are depicted in **Figure S5**, which shows that salt promotes the formation of a single dense aggregate, whereas the salt-free system forms smaller, structurally heterogeneous clusters.

### Energetic signatures of aggregation: electrostatics and van der Waals contributions

To assess the thermodynamic favourability of the observed aggregation, electrostatic and van der Waals (vdW) energy profiles were computed over the simulation trajectories for both systems (**Figure 6 d**). The salt-containing system, which formed a large and stable aggregate early in the simulation, exhibited a rapid and substantial decrease in electrostatic energy, reflecting enhanced inter-peptide charge screening and favourable surfactant–peptide interactions. This sharp drop in electrostatic energy is indicative of aggregation-driven stabilisation, facilitated by the presence of salt, which diminishes repulsive interactions among the positively charged peptide residues.

In comparison, the salt-free system showed a more gradual decline in electrostatic energy over time. Although the overall trend eventually approached a similar energy minimum, the slower kinetics reflect delayed aggregation due to the absence of ionic screening, which maintains stronger electrostatic repulsion between peptides in the early phase of the simulation.

A parallel trend was observed in van der Waals energies, where both systems exhibited a consistent reduction over time. In particular, the salt-containing system showed an earlier and steeper decline, corresponding to the tight packing of peptides within a large aggregate. The salt-free system followed a similar path with a lag, as clustering proceeded more slowly and less cooperatively. The reduction in van der Waals energy across both systems supports the occurrence of aggregation events, as closer packing of hydrophobic surfaces leads to energetically favourable dispersion interactions. Different energetic decompositions over time (**Figure S6**) show that salt-free systems exhibit stronger electrostatic and van der Waals stabilisation, consistent with their tighter and more dynamic aggregation.

These analyses collectively suggest that SDS at below-CMC concentrations can facilitate peptide aggregation under crowded conditions, particularly when the surfactant-to-peptide ratio is low, and salt is present. Rather than acting purely as solubilising agents, SDS monomers in these contexts appear to behave as aggregation-promoting scaffolds, enabling the clustering of peptides by electrostatically screening repulsive interactions, particularly in the presence of salt; providing localized hydrophobic environments that promote peptide association and stabilizing early aggregation nuclei, particularly via N-terminal β-sheet formation. The convergence of contact map signatures and energetic profiles in both salt and salt-free systems further underscores that SDS monomers below CMC can drive aggregation through distinct mechanisms, dictated by the surrounding ionic and concentration environment. The salt-containing system achieves aggregation more rapidly and cooperatively, while the salt-free system does so more gradually, with multiple intermediate cluster states.

To elucidate the role of ionic strength in modulating the aggregation behaviour of Uperin 3.5, we conducted comparative all-atom molecular dynamics simulations of two peptide–SDS systems, differing solely in the presence or absence of physiological salt (0.15 M NaCl). In both setups, the simulations were performed under below-CMC conditions for sodium dodecyl sulphate (SDS), ensuring the surfactant existed as dissociated monomers rather than as micelles. In Figure 7, the peptide ensemble in the presence of 0.15 M NaCl, comprising 20 Uperin 3.5 peptides—including two preformed β-sheet tetramers and 12 disordered monomers—along with 12 SDS monomers, evolves over 1 μs into a single, compact aggregate. This heterogeneous assembly incorporates multiple structural motifs, including α-helices, β-sheets, and unstructured peptides, all closely associated with the SDS monomers. The presence of salt plays a critical role in reducing electrostatic repulsion among the predominantly cationic peptides, thereby stabilizing peptide–peptide and peptide–SDS interactions, enhancing structural packing, and promoting large-scale aggregation. In contrast, under salt-free conditions, an equivalent peptide ensemble with 12 SDS monomers exhibits markedly reduced aggregation. Rather than forming a unified structure, the peptides organize into multiple smaller clusters of 5–6 peptides, which remain spatially distinct and display varied conformational outcomes, including α-helical, β-sheet, or mixed secondary structures. The absence of ionic screening likely preserves inter-peptide electrostatic repulsion, thereby limiting long-range assembly even in the presence of SDS. To quantify the relationship between structural association and energetic stabilization, regression analysis was performed correlating the number of inter-peptide contacts with total interaction energy. A higher correlation coefficient (R² = 0.63) is observed under salt-free conditions compared to the salt-containing case (R² = 0.53), indicating that in smaller, more ordered clusters, interaction energy scales more linearly with the number of contacts. In contrast, the large and structurally heterogeneous aggregate formed in the presence of salt likely involves additional stabilizing contributions from side-chain packing, solvent-mediated effects, and non-canonical interactions, resulting in a less direct relationship between contact number and energy.

**Figure 7.**
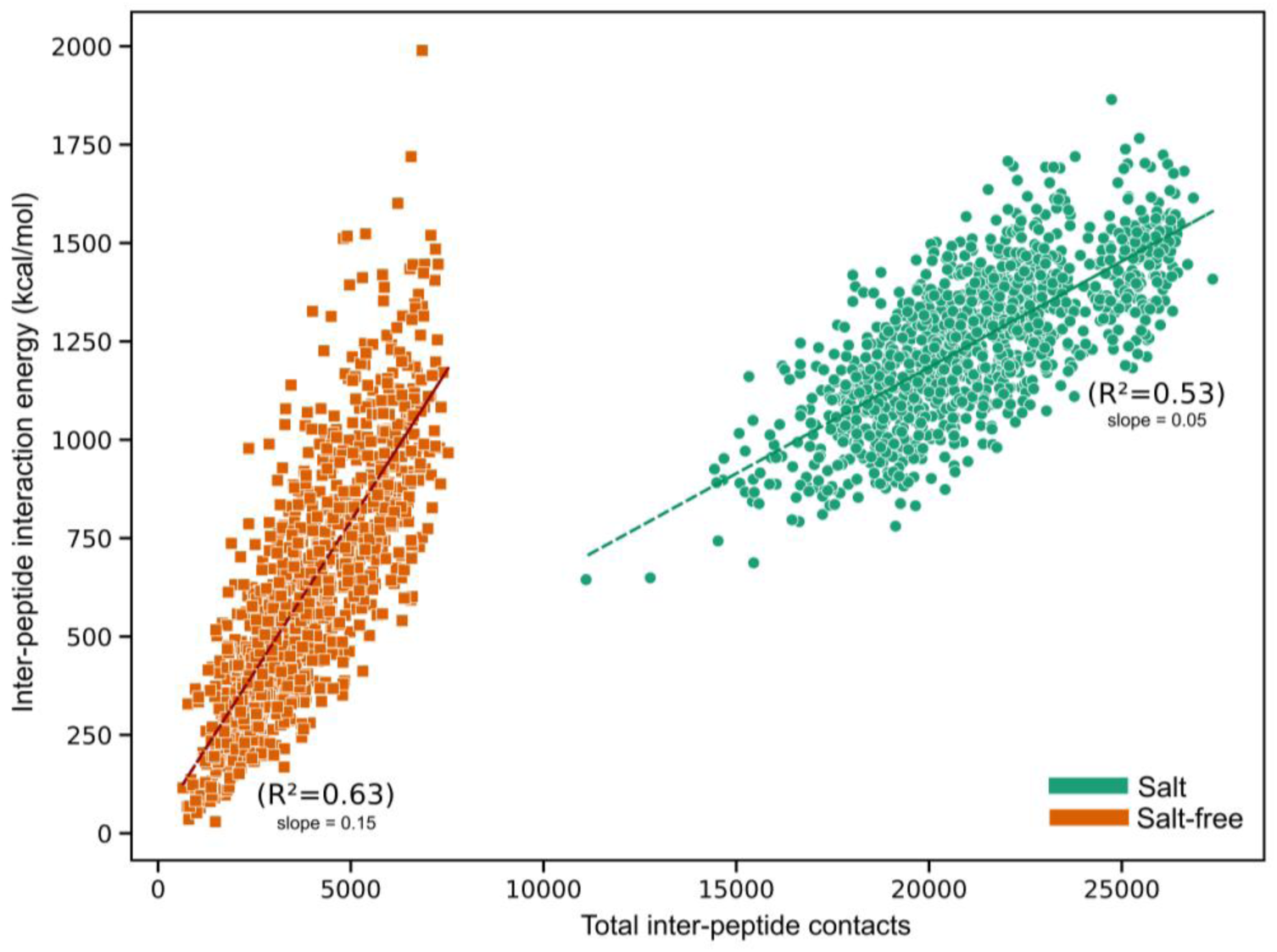
Correlation between inter-peptide contact number and peptide–peptide interaction energy in Uperin 3.5–SDS simulations with and without salt. Each data point represents an individual peptide under two conditions: in the presence and absence of 0.15 M NaCl. In the presence of salt, a peptide ensemble comprising 20 peptides (two preformed β-sheet tetramers and 12 random coils) and 12 dissociated SDS monomers forms a single large heterogeneous aggregate during the 1 μs simulation. This aggregate contains both β-sheet and α-helical structures stabilized around the SDS monomers. In contrast, under salt-free conditions, 20 initially random coil peptides with 12 SDS monomers remain dispersed as several smaller clusters of 5–6 peptides, with secondary structures forming locally. Linear regression analysis reveals a moderate correlation (R² = 0.53) in the presence of salt, reflecting the structural diversity of the large aggregate and a broader range of interaction energies. In the absence of salt, a stronger correlation (R² = 0.63) is observed, indicating that interaction energy scales more predictably with inter-peptide contacts in smaller, more homogeneous clusters. These results demonstrate that ionic strength significantly modulates peptide self-assembly dynamics and structural organization.

Collectively, these findings demonstrate that SDS monomers below CMC can facilitate peptide aggregation, but the extent and nature of the assembly are strongly influenced by ionic strength. The presence of salt significantly enhances aggregation propensity by screening electrostatic repulsion and promoting both compact packing and conformational transitions, particularly the emergence of β-sheet and α-helical structures. These results underscore the importance of environmental factors such as ionic strength and surfactant identity in regulating peptide self-assembly and support the concept of environment-responsive functional amyloidogenesis in amphipathic antimicrobial peptides like Uperin 3.5.

### Surfactant environments drive a structural switch between amyloid and antimicrobial states

To experimentally test the simulation-predicted dependence of Uperin 3.5 structure on surfactant identity and concentration, circular dichroism (CD) spectroscopy and thioflavin-T (ThT) fluorescence assays were performed under a range of membrane-mimetic conditions. These complementary measurements simultaneously monitor secondary-structure evolution and amyloid formation as the peptide transitions between disordered, α-helical, and β-sheet-rich states. By systematically varying surfactant concentration relative to the critical micelle concentration (CMC), the experiments directly probe how amphiphilic environments regulate peptide self-assembly.

As shown in **Figure 8**, surfactant environments strongly modulate the aggregation behaviour of Uperin 3.5. In aqueous solution, the peptide exhibits measurable amyloid formation following the addition of PBS buffer, as indicated by increased ThT fluorescence. In contrast, micellar concentrations of surfactant markedly suppress aggregation. In particular, SDS above the CMC (10 and 50 mM) and DPC across the concentrations tested display minimal ThT fluorescence, indicating that peptide self-assembly is strongly inhibited in micellar environments.

**Figure 8.**
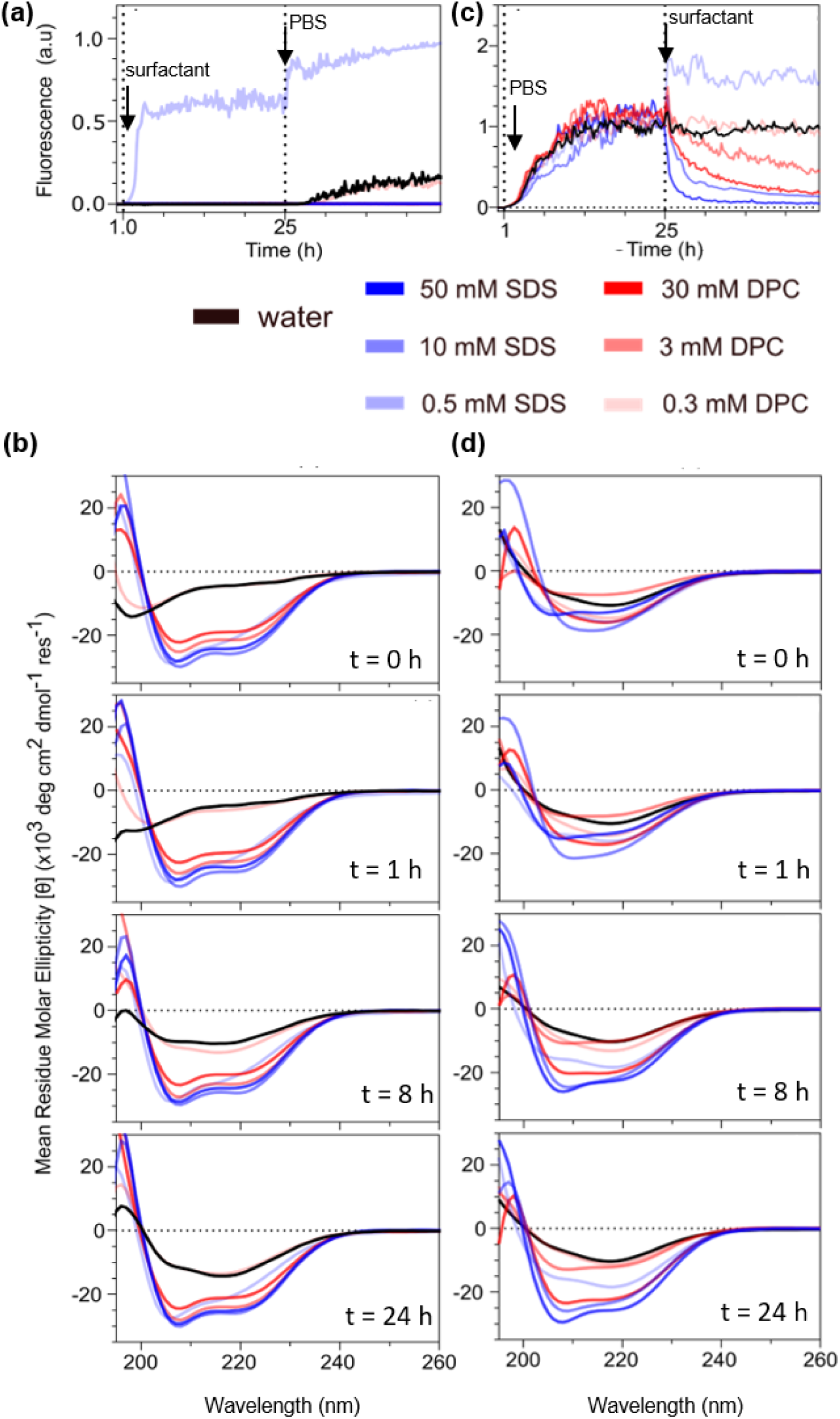
Effect of surfactant concentration on Uperin 3.5 aggregation and secondary structure (a) Aggregation of Uperin 3.5 in the presence of surfactants. ThT fluorescence kinetics of Uperin 3.5 (125 μM) were recorded after addition of surfactants followed by PBS to initiate aggregation. Fluorescence intensities are normalized to the maximum signal. (b) Corresponding circular dichroism (CD) spectra recorded immediately after PBS addition and at 1, 8, and 24 h show that micellar concentrations of SDS (10 and 50 mM) and DPC stabilize α-helical conformations (minima at ∼208 and 222 nm), whereas below-CMC SDS (0.5 mM) promotes β-sheet formation and enhanced aggregation**(c)** Effect of surfactants on pre-aggregated peptide. Uperin 3.5 was first allowed to aggregate in PBS, after which surfactants were added. ThT fluorescence kinetics are normalized to the fluorescence intensity prior to surfactant addition. (d) CD spectra recorded at the same time points show that micellar surfactants promote conversion of aggregated peptide into α-helical conformations, whereas below-CMC SDS maintains β-sheet characteristics.

Circular dichroism spectra reveal that this suppression of aggregation coincides with stabilization of α-helical conformations. Under micellar conditions, the peptide adopts a characteristic amphipathic helix, evidenced by the canonical double minima at approximately 208 and 222 nm. Such helices arise from electrostatic attraction between surfactant headgroups and the cationic residues of Uperin 3.5, followed by hydrophobic insertion of non-polar side chains into the micellar environment. This interaction stabilizes the peptide in a monomeric or loosely associated helical state and prevents the intermolecular β-sheet interactions required for amyloid nucleation.

A distinct behaviour is observed under below-CMC SDS conditions (0.5 mM). Despite the appearance of helical features in the CD spectra, ThT fluorescence simultaneously indicates enhanced aggregation following salt addition. This suggests that SDS monomers partially screen electrostatic repulsion between the positively charged residues of Uperin 3.5, facilitating peptide association while still permitting transient helical structure. Such behaviour is consistent with the formation of intermediate or non-canonical assemblies that precede β-sheet nucleation. In contrast, below-CMC DPC (0.3 mM) produces minimal structural change and exhibits aggregation behaviour similar to peptide in water, reflecting weaker electrostatic interactions between the zwitterionic headgroup and the cationic peptide.

To determine whether surfactant environments can actively remodel existing peptide aggregates, a complementary experimental protocol was performed in which Uperin 3.5 was first allowed to self-assemble in PBS prior to surfactant addition (**Figure 8c**). In this case, the peptide initially displays the characteristic sigmoidal ThT fluorescence profile of amyloid formation. Introduction of micellar surfactants results in rapid and concentration-dependent disassembly of these aggregates. Addition of 50 mM SDS produces an immediate decrease in ThT fluorescence, while 10 mM SDS and DPC micelles induce a slower but comparable reduction, demonstrating that the extent and rate of disaggregation scale with surfactant concentration and interaction strength.

These fluorescence changes are accompanied by a corresponding structural transition in the CD spectra (**Figure 8d**). Following surfactant addition, the β-sheet signature near 218 nm progressively diminishes, while the α-helical minima at 208 and 222 nm emerge. This observation indicates that micellar surfactants not only solubilize peptide aggregates but actively remodel their secondary structure toward amphipathic helical conformations.

Consistent with the aggregation experiments described above, below-CMC SDS again exhibits anomalous behaviour. Rather than dissolving aggregates, 0.5 mM SDS enhances ThT fluorescence and strengthens β-sheet signatures in the CD spectra, indicating that partial charge screening by SDS monomers promotes additional peptide association.

Taken together, these results reveal a clear dependence of peptide structural fate on surfactant charge and concentration. The anionic surfactant SDS induces α-helicity more efficiently than the zwitterionic DPC, reflecting stronger electrostatic attraction between negatively charged sulfate headgroups and the cationic residues of Uperin 3.5. Once micelles form, this interaction promotes peptide solubilization and stabilizes amphipathic α-helical conformations that suppress β-sheet stacking and amyloid growth. In contrast, below-CMC SDS conditions permit partial charge screening without providing a stabilizing micellar interface, thereby facilitating peptide association and β-sheet nucleation.

Overall, the combined CD and ThT measurements provide strong experimental validation of the simulation predictions. Micellar surfactant environments stabilize α-helical conformations and disrupt amyloid assemblies, whereas below-CMC SDS uniquely promotes β-sheet formation. These findings highlight an environmentally regulated structural switch in Uperin 3.5, whereby membrane-mimetic environments bias the peptide away from amyloidogenic aggregation and toward biologically relevant antimicrobial conformations.

As a summary of these results, **Figure 10** integrates structural outcomes across all surfactant regimes explored in this study. The conceptual landscape highlights how surfactant charge, concentration relative to the CMC, ionic strength, and peptide stoichiometry cooperatively determine whether Uperin 3.5 adopts α-helical antimicrobial conformations or β-sheet-rich amyloid assemblies. This emphasises that the AMP to Amyloid transition is not an intrinsic property of the peptide, but rather an emergent, environment-dependent response. The detailed relationship between α/β structural distributions and the underlying free-energy landscape is further illustrated in the two-dimensional phase maps in **Figure S7**, which compare all six SDS surfactant environments. An integrative summary of surfactant charge, assembly state, and their effects on Uperin 3.5 structure and aggregation is provided in **Supplementary Table ST3**.

**Figure 10.**
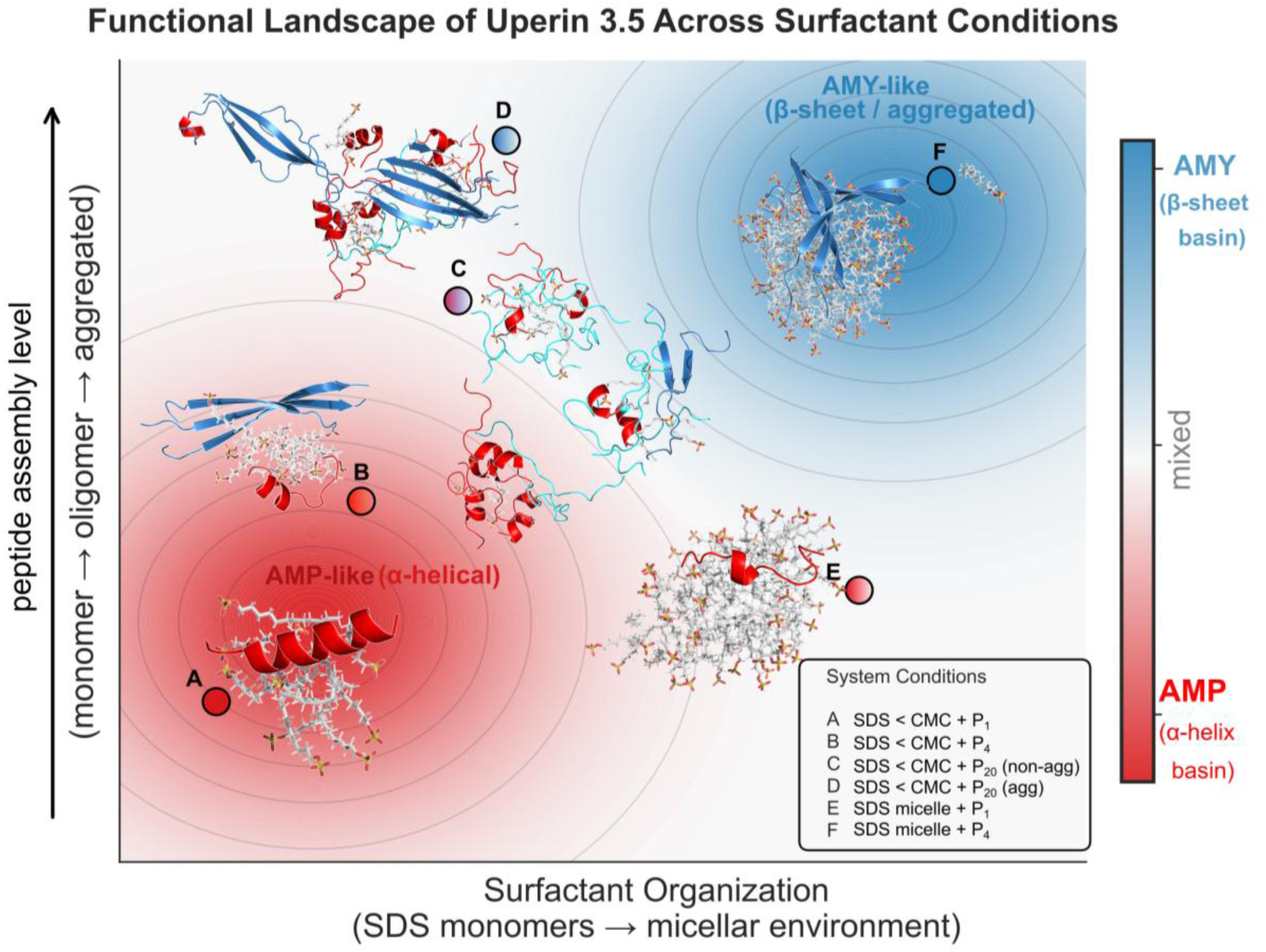
Functional Switching Landscape of Uperin 3.5 Across Surfactant Conditions. A unified structural map summarising how surfactant identity, concentration relative to the CMC, peptide crowding, and ionic strength shape the conformational fate of Uperin 3.5. Anionic SDS strongly promotes α-helical antimicrobial states at above-CMC concentrations, while zwitterionic DPC induces moderate helicity. Under below-CMC conditions, limited SDS availability and peptide crowding favour β-sheet-rich amyloid states, further enhanced by salt. Above the CMC, micellar interfaces solubilise peptides and suppress β-sheet formation, promoting antimicrobial α-helical conformations. Collectively, the landscape illustrates an environmentally guided AMP to Amyloid switch governed by surfactant assembly dynamics, stoichiometry, and ionic strength.

## Conclusions

The combined experimental and simulation studies revealed that both the identity and concentration of surfactants play crucial roles in determining the peptide’s structural fate. At high-CMC concentrations, SDS and DPC stabilise α-helical conformations and suppress β-sheet aggregation, with the effect being more pronounced for the anionic SDS due to stronger electrostatic interactions with cationic peptide residues. In contrast, below-CMC SDS conditions uniquely promote aggregation, driving peptides toward mixed α/β conformations and enhancing β-sheet clustering. Salt further accelerates this process by screening electrostatic repulsion between peptides, thereby lowering the barrier for aggregation and favouring amyloid formation. Together, these results highlight a finely tuned balance: high surfactant-to-peptide ratios bias peptides toward antimicrobial helices, while limited solubilising capacity or ionic screening promotes amyloidogenic assembly. Importantly, the convergence of CD, ThT, and simulation data validates a functional switching mechanism, whereby antimicrobial peptides can reversibly transition between helical and amyloid states depending on environmental context. This AMP ↔ Amyloid switch is therefore dictated by surfactant identity, concentration, and ionic conditions, underscoring the adaptive plasticity of these peptides and offering mechanistic insights into their dual functional roles.

## Acknowledgements

The authors acknowledge financial support from the Department of Biotechnology (DBT), Government of India (GOI), for the project (IMURA0956) at IITB-Monash Research Academy and for the award of a PhD stipend to SB. DC acknowledges Monash University for the award of an RTP stipend PhD scholarship. DC add your scholarship details….

## Supplementary Information

### Experimental methods

#### Materials

U3.5 peptide, amidated at the C-terminus, was purchased from Peptide 2.0 Inc. (98% purity, Chantilly, VA). Potassium phosphate monobasic (KH_2_PO_4_, anhydrous 99%) and potassium phosphate dibasic (K_2_HPO_4_, anhydrous), purchased from Sigma-Aldrich (St. Louis, USA), were used for the preparation of phosphate-buffered saline (PBS) solution. Sodium chloride (NaCl ≥ 99.5%) was used for preparation of PBS solution. Ultrapure water (18.2 MΩ cm) was used for buffer preparation and peptide dissolution in all experiments. In all the experiments, 10-fold concentrated PBS was used to make a final concentration of 20 mM phosphate and 100 mM NaCl (1 ×PBS). The thioflavin T (ThT, Sigma-Aldrich) was diluted in dimethyl sulfoxide (DMSO) to make a 1 mM stock solution. The ThT stock was protected from light and stored at -20 °C. The final concentration of ThT in the assay was 10 µM. For experiments involving surfactants, sodium dodecyl sulfate (SDS) and dodecyl phosphocholine (DPC) were purchased from Pall Life Sciences, New York, USA.

#### Preparation of peptide stocks

500 µM stock solutions of Uperin 3.5 were prepared in ultrapure water in 1 mL aliquots and stored in low-binding Eppendorf tubes sealed with parafilm. All peptide powder and stocks were stored at -20°C until required.

#### Buffer preparation

A phosphate-buffered saline (PBS) solution comprised of 20 mM phosphate and 100 mM NaCl was used in fluorescence and circular dichroism studies. The PBS solution was prepared as a ten-fold concentrated (10x PBS) to allow for addition of species to the required dilution for the desired final concentration. The phosphate buffer was prepared in approximately a 2:1 ratio of K_2_HPO_4_ to KH_2_PO_4_. The pH of the PBS solution was then adjusted to 7.40 ± 0.02 using NaOH and HCl solutions where necessary. The PBS buffer solution was filtered through a 0.2 µM hydrophilic polypropylene membrane filter (Pall Life Sciences, New York, USA) and stored at 4°C for a maximum of two weeks.

#### Preparation of surfactant solutions

Sodium dodecyl sulfate (SDS) was prepared as a 500 mM stock solution in water and filtered through a 0.2 µM membrane filter. SDS stock solutions were stored at room temperature (∼ 21 C) for a maximum of 2 weeks. Where intended, SDS was used at 50 mM and 10 mM in Circular Dichroism (CD) spectroscopy and fluorescence assays (using ThT) to ensure SDS micelles were formed. Some studies were conducted below the critical micelle concentration, using 0.5 mM SDS solution. Dodecylphosphocholine (DPC) were prepared as either a 300 mM or 30 mM stock solution in 100 µL aliquots of chloroform, which was evaporated under N_2_ and then resuspended in pure water to achieve the desired concentration. DPC stock solutions were stored at -20°C until required. DPC was used at 30 mM and 3 mM in Circular Dichroism spectroscopy and fluorescence assays to ensure micelles were formed. For experiments that required sub-micellar concentrations, 0.3 mM DPC was used.

#### Circular Dichroism

The peptide stock solutions were thawed to room temperature and diluted to a concentration of 125 µM in water, such that following the introduction of high salt PBS buffer (10% v/v) and surfactant (10% v/v) to the final concentration of 100 µM. The typical CD methodology was to first measure the spectrum of the peptide in pure water, prior to the introduction of buffer. Measurements in water were conducted at 10 min intervals for an hour, and hourly for 24 hours following the addition of buffer. Finally, surfactant was introduced and measured every hour for 24 hours. All CD measurements were recorded with a Jasco J-815 CD spectropolarimeter using a 1.0 mm path length quartz cuvette. The temperature of the cuvette holder was controlled using a water jacket and set to 37° C throughout the experiment. Continuous scan mode was used and a 2.0 nm bandwidth with 100 nm/min scan rates and a 2 s response time. CD measurements were recorded from 550 nm to 190 nm if in water, or to 195 nm if PBS buffer was present due to interference by chloride ions at wavelengths below 195 nm. For time-course experiments, where hourly measurements were required over 24 hours or longer duration, an automatic timer (AutoClicker, Polar) was set to activate the spectropolarimeter at hourly intervals over the course of the measurement.

The data was processed using the JASCO Spectra Manager software (version 2). Baselines were recorded prior to each experiment and subtracted from those spectra containing peptides. The baseline corrected spectra were smoothed using binomial smoothing with 5 iterations. The data were transformed from millidegrees (*θ_λ_*) to mean residue molar ellipticity (*θ*) to allow for data comparisons between different concentrations by using Equation 1. CD data were plotted using GraphPad PRISM 9 software.

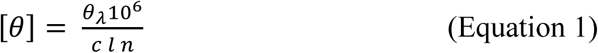

Where [*θ*] is the mean residue molar ellipticity, *θ_λ_* is the measured ellipticity in millidegrees, *c* is the concentration in µM, *l* is the pathlength in mm, and *n* is the number of peptide bonds (16 for U3.5 peptides).

#### ThT fluorescence self-assembly assay

Thioflavin T (ThT) was prepared as a 1.0 mM stock solution by dissolution in DMSO. It was stored at -20°C and protected from light using aluminum foil. For the ThT fluorescence assays, black polystyrene 96-well microplates with bottom optic, non-binding surfaces (Greiner Bio-One, Germany) were used. Fluorescence data were obtained using a CLARIOstar plate reader (BMG Labtech, Germany) with excitation and emission wavelengths of 440-10 and 480-10 nm, respectively. All experiments were performed at 37 °C and measurements were recorded every 10 minutes for the duration of each experiment. To prepare the microplate wells for triplicates, peptide stock and water were combined in a non-binding Eppendorf with 1 mM ThT. These solutions were vortexed and immediately transferred into the wells. The microplate was sealed using a ThinSeal^TM^ adhesive film (Austral Scientific, Australia) to prevent sample evaporation. After the measurement of the peptides in water were completed, the adhesive seal was temporarily removed to allow for the addition of PBS (10% v/v) and/or surfactant, respectively, into each well to initiate peptide aggregation.

Baselines were subtracted from the raw data using MARS data analysis software (BMG, Labtech, version no.?). These data were analyzed following visual inspection of each replicate and graphing the average data using GraphPad PRISM 9 (version no.?).

### Simulation methods

All-atom molecular dynamics (MD) simulations were performed to investigate the interactions between Uperin 3.5 (U3.5) peptides—both as monomers and tetramers—and three surfactants of differing charge characteristics: anionic sodium dodecyl sulphate (SDS), zwitterionic dodecyl phosphocholine (DPC), and cationic cetyltrimethylammonium bromide (CTAB)^1–4^. Simulations were conducted under two principal conditions: (i) below the critical micelle concentration (CMC), where individual surfactant monomers were randomly distributed around the peptide(s) and (ii) above the CMC, where preformed micelles were introduced into the system, and peptides were placed in proximity to the micellar surface.

Initial system configurations were generated using Packmol^5^, ensuring random, non-overlapping placement of surfactants and peptides. For micelle-based systems, spherical micelles composed of 60 SDS^6,7^ or DPC molecules (∼40 Å diameter)^8^ and 120 CTAB ^9^molecules (∼50 Å diameter) were constructed in accordance with experimentally reported micelle sizes. The micelle centre of mass was harmonically restrained at the centre of the simulation box to maintain micelle position throughout the simulation.

All simulations were carried out using the NAMD 2.14^10^ software package with the CHARMM36m^11^ force field for peptides and surfactants, and the TIP3P water model^12^. Systems were solvated in explicit water with 0.15 M NaCl unless otherwise specified. Simulation boxes were orthorhombic and subjected to periodic boundary conditions in all directions. Long-range electrostatics were computed using the particle mesh Ewald (PME) method^13^ with a grid spacing of 1 Å. Lennard-Jones interactions were smoothed using a force-switching function between 10 and 12 Å. Temperature and pressure were maintained using a Langevin thermostat and Nosé–Hoover Langevin barostat at 310 K and 1 atm, respectively.

Peptides were initially restrained using a harmonic potential of 10 kcal mol⁻¹ Å⁻² to permit solvent equilibration, with restraints gradually released over 2 ns. This was followed by a 400 ps energy minimisation and a two-stage equilibration process: 1 ns under NVT conditions and 1 ns under NPT conditions. Production runs were then performed in the NPT ensemble (T = 310 K, P = 1 atm) using a 2 fs integration timestep.

### Peptide preparation and system categories

Monomeric U3.5 peptides with a variety of secondary structures were extracted from long-timescale (1–2 μs) simulations of single or multiple peptides in water and saline (0.15 M NaCl) environments, yielding initial conformations with dominant random coil or partial α-helical character. The C-terminal was amidated, in accordance with experimental data. Tetrameric peptide structures were taken from the cryo-EM structure (PDB ID: 7QV5), which displays a parallel β-sheet propeller-like arrangement.ref needed here.

The categories of systems that were studied:

1. **Sub-CMC simulations:** Monomeric or tetrameric peptides were surrounded by surfactant monomers: 12 SDS, 8 DPC or 8 CTAB molecules, each randomly dispersed around the peptides using PACKMOL. During equilibration, the surfactant centres of mass were constrained and subsequently released during production runs to allow free peptide–surfactant interactions.
2. **Micelle-based simulations:** Monomeric or tetrameric peptides were placed near the surface of preformed micelles of SDS, DPC or CTAB. The micelle was held fixed during simulation via harmonic restraints on its centre of mass, while peptides interacted freely with the micellar surface.
3. **3. 20-Peptide simulation with or without salt**

To probe aggregation under physiologically relevant crowding conditions, two 20-peptide systems were constructed:

- A non-aggregated system consisting of 20 peptides in random coils and 12 SDS monomers (sub-CMC), simulated in the absence of any salt (i.e. no NaCl).
- An aggregated system comprising 2 preformed tetramers and 12 peptides (totalling 20 peptides) were placed randomly in the simulation box, along with 12 SDS monomers and salt (0.15 M NaCl).

Both these setups followed the same equilibration protocol as above, with initial harmonic restraints applied only during the solvent relaxation phase. These simulations allowed for the assessment of salt-dependence of aggregation behaviour in the presence of sub-CMC SDS, consistent with data obtained in the experimental ThT assays. The simulation setup details have been provided in the Tables ST1 and ST2 below, respectively.

---

**Table ST1:** Detailed information of simulation setup.

| Surfactant Simulations | SDS | DPC | CTAB |
| --- | --- | --- | --- |
| Monomer simulations<br>(Below CMC) |  |  |  |
| Peptides No.=1 |  |  |  |
| Surfactant Molecules | 12 | 8 | 8 |
| Box size ($\text{\AA}^3$) | 120 | 120 | 120 |
| Total atoms | 166317 | 166489 | 166502 |
| Simulation time (ns) | 1000 | 1000 | 1000 |
| Monomer simulations<br>(Above CMC) |  |  |  |
| Peptides No.=1 |  |  |  |
| Surfactant Molecules | 60 | 60 | 120 |
| Box size ($\text{\AA}^3$) | 120 | 120 | 120 |
| Total atoms | 115296 | 114782 | 166530 |
| Simulation time (ns) | 1000 | 1000 | 1000 |
| Tetramer simulations<br>(Below CMC) |  |  |  |
| Peptides No.=4 |  |  |  |
| Surfactant Molecules | 12 | 8 | 8 |
| Box size ($\text{\AA}^3$) | 120 | 120 | 120 |
| Total atoms | 166125 | 166172 | 166256 |
| Simulation time (ns) | 1000 | 1000 | 1000 |
| Tetramer simulations<br>(Above CMC) |  |  |  |
| Peptides No.=4 |  |  |  |
| Surfactant Molecules | 60 | 60 | 120 |
| Box size ($\text{\AA}^3$) | 120 | 120 | 120 |
| Total atoms | 115602 | 114681 | 166408 |
| Simulation time (ns) | 1000 | 1000 | 1000 |

**Table ST2:**
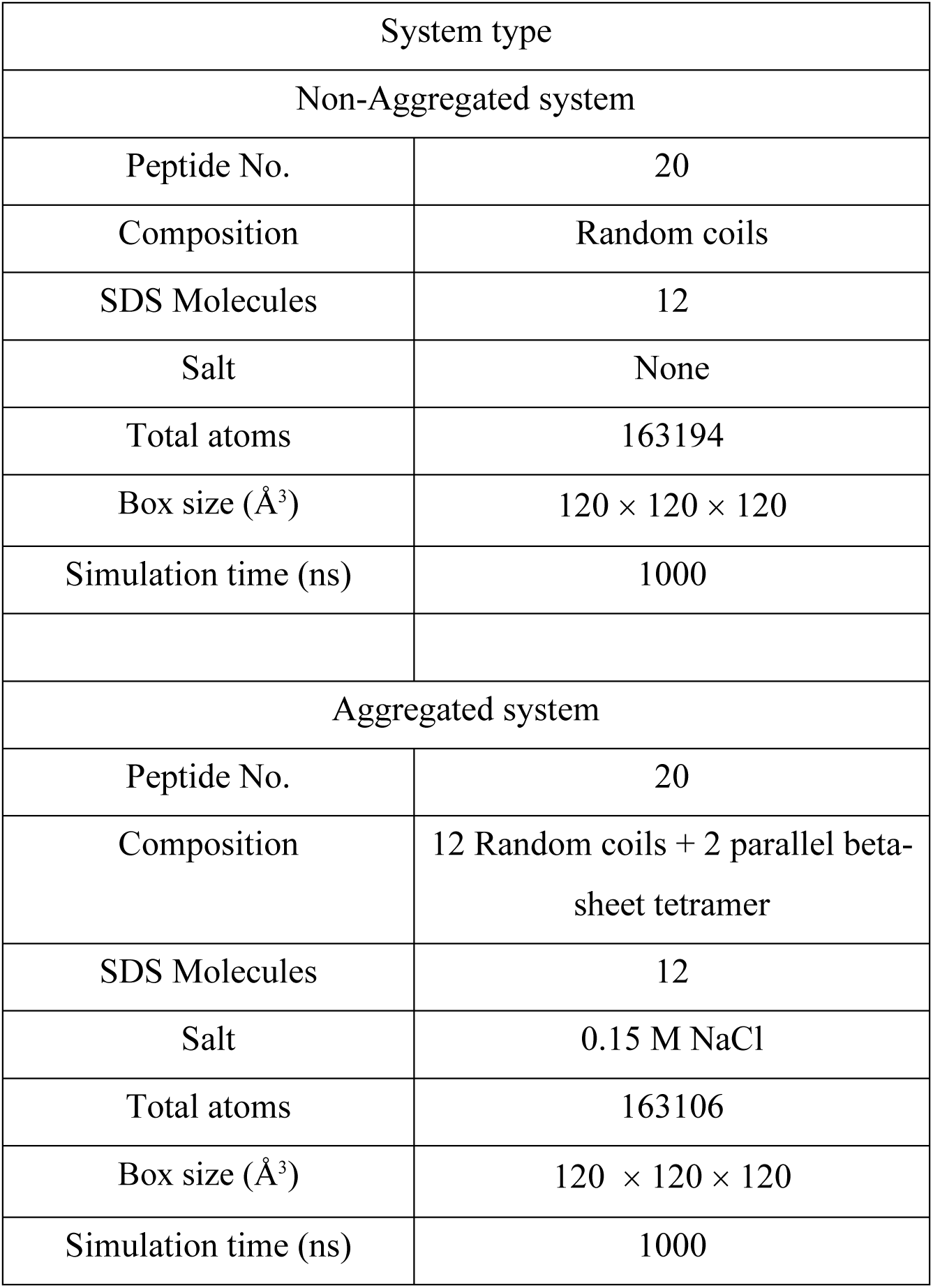
Detailed information of simulation setup.

**Table ST3:**
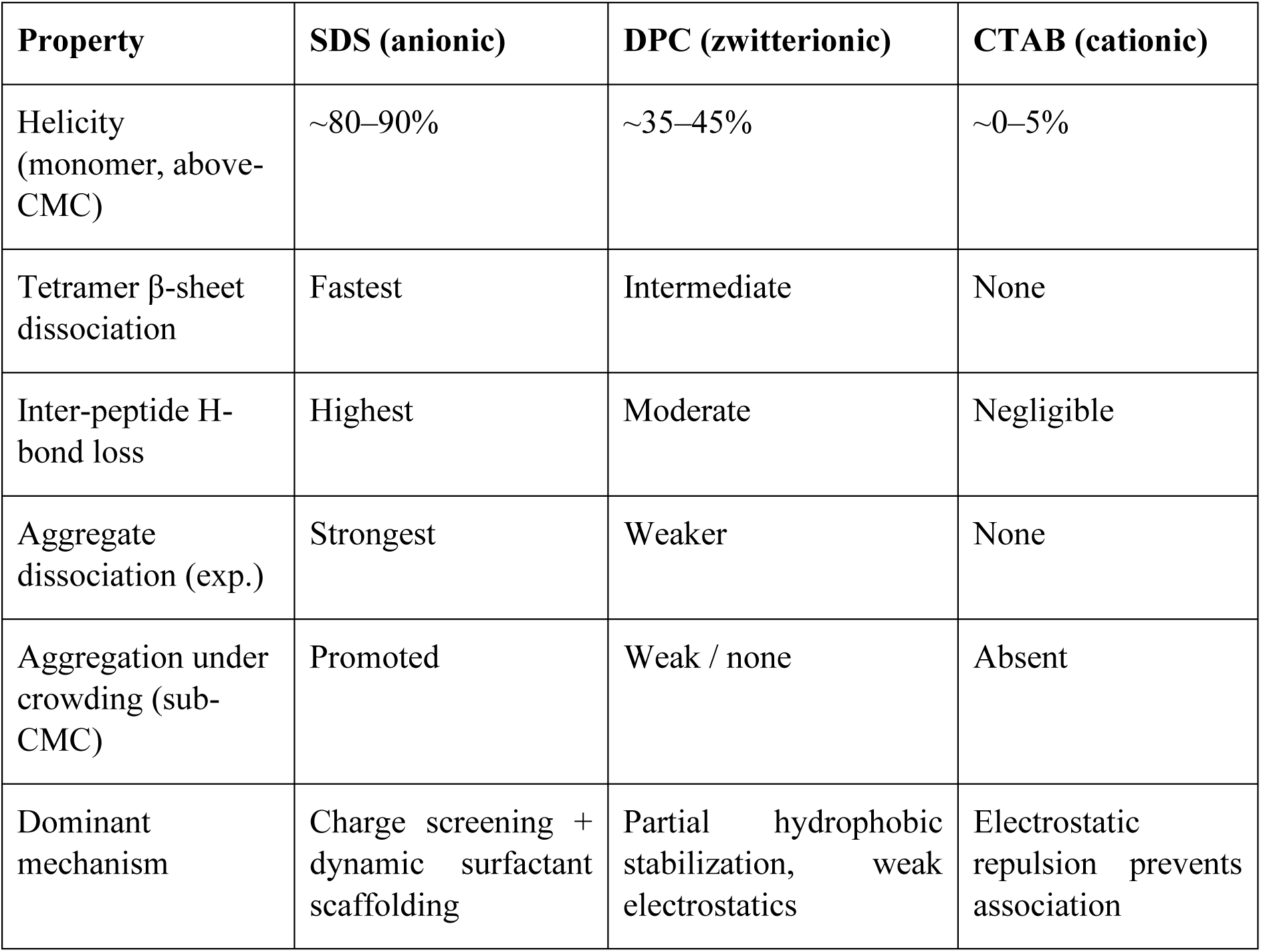
Summary of U3.5 peptide behaviour with surfactants.

**Figure S1.**
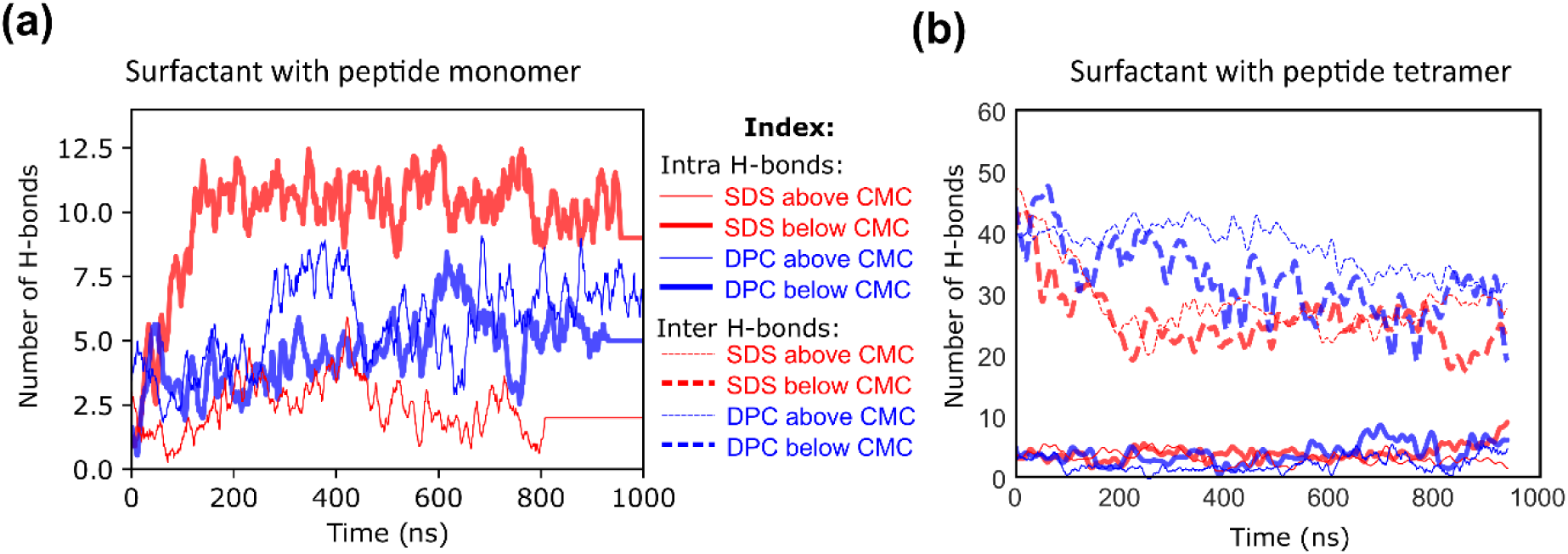
(a)Number of intramolecular hydrogen bonds (H-bonds) formed by the peptide monomer in the presence of SDS and DPC, at surfactant concentrations below and above the critical micelle concentration (CMC) over the simulation trajectory. (b) Variation in intra- and intermolecular hydrogen bond formation within the peptide tetramer upon interaction with SDS and DPC under sub-CMC and above-CMC conditions along the simulation trajectory.

**Figure S2.**
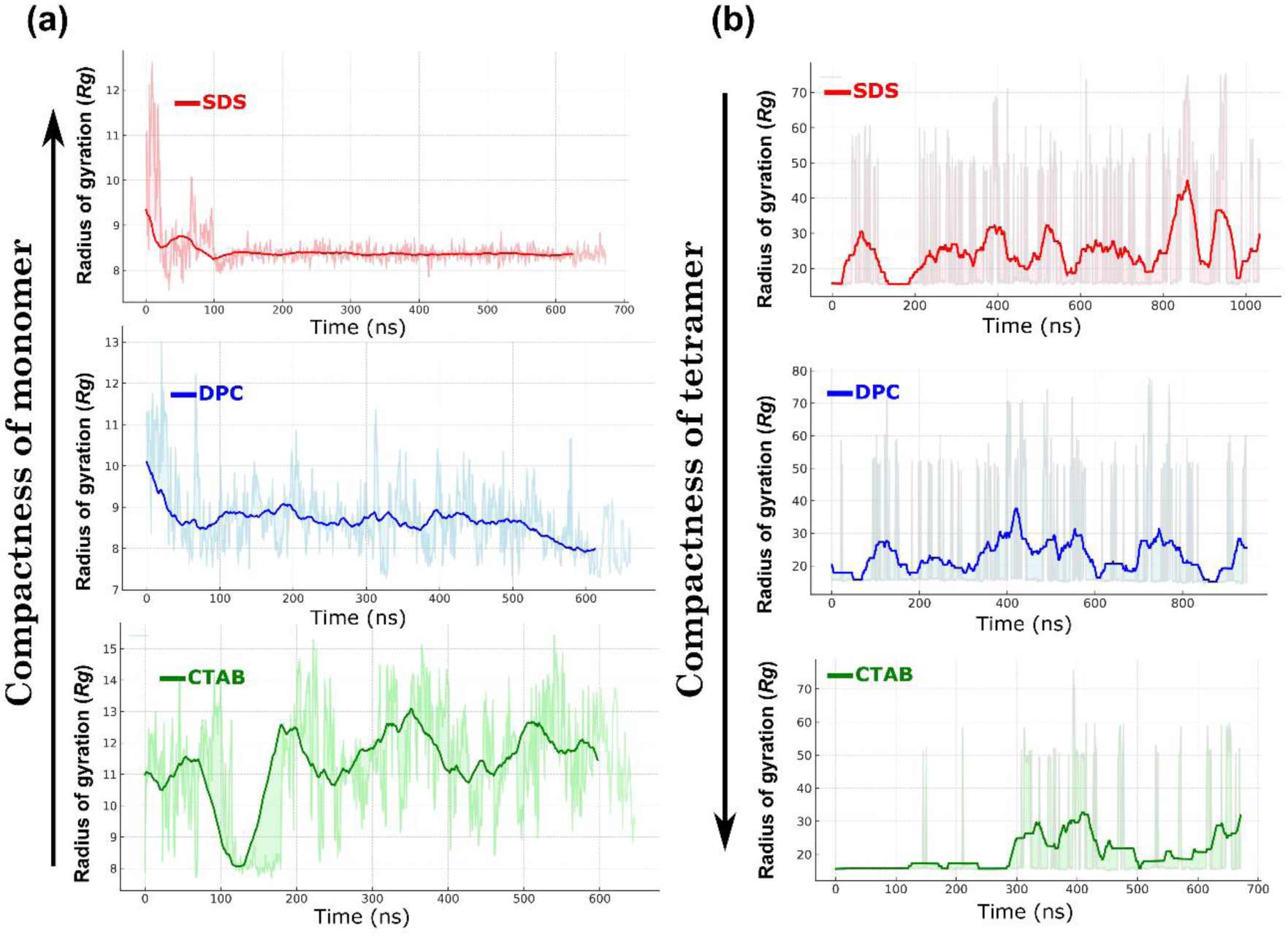
(a) Time evolution of the radius of gyration (*Rg*) for the peptide monomer in the presence of surfactants with different head group charges—SDS (anionic), DPC (zwitterionic), and CTAB (cationic). The monomer exhibits the least fluctuations in *Rg* with SDS, indicating rapid stabilization into a compact helical conformation. Intermediate fluctuations are observed with DPC, where partial helix formation (approximately 40–50%) occurs, while in CTAB, the peptide shows minimal interaction and remains largely unstructured. (b) In contrast, *Rg* variations for the peptide tetramer reveal an inverse trend. SDS induces the highest fluctuations due to disruption of inter-peptide hydrogen bonds, likely driven by strong electrostatic interactions, resulting in reduced compactness. DPC shows moderate fluctuations, while the tetramer remains largely stable in CTAB, suggesting limited disruption of the oligomeric structure.

**Figure S3.**
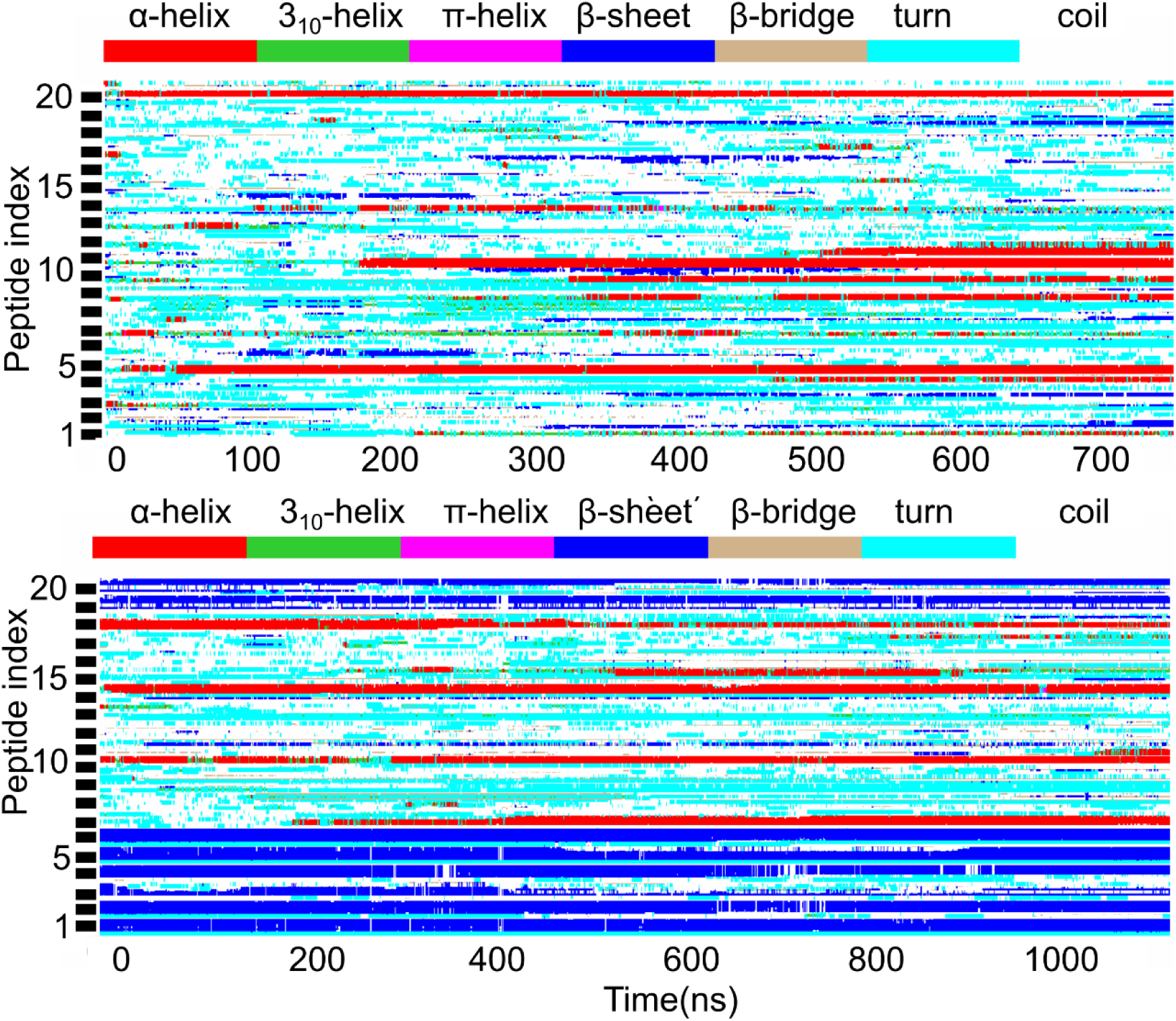
Heat maps illustrating the time-resolved secondary structure evolution for individual peptides in both in Uperin 3.5 peptide–SDS systems in the absence (a) and in the presence (b) of salt, color-coded by secondary structure type (α-helix, β-sheet, or coil).

**Figure S4.**
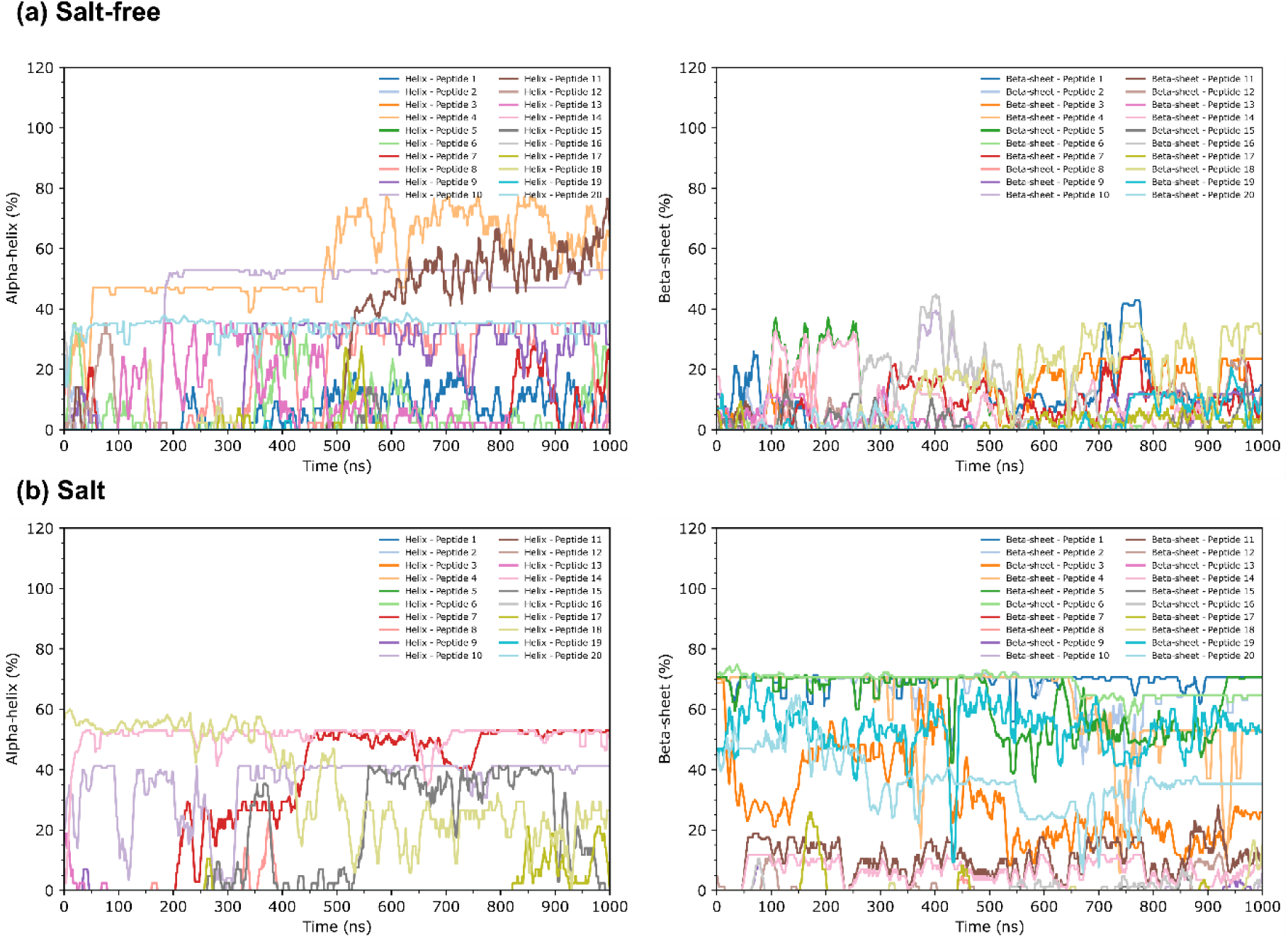
Percentage evolution of secondary structure over time for each individual peptide in the 20-peptide system under different conditions. (a) System comprising 20 initially unstructured peptides (random coils) and 12 SDS monomers, in the absence of salt and under sub-CMC conditions. (b) System comprising two pre-formed β-sheet tetramers, 12 peptides (random coil), and 12 SDS monomers, in the presence of salt and under sub-CMC conditions.

**Figure S5.**
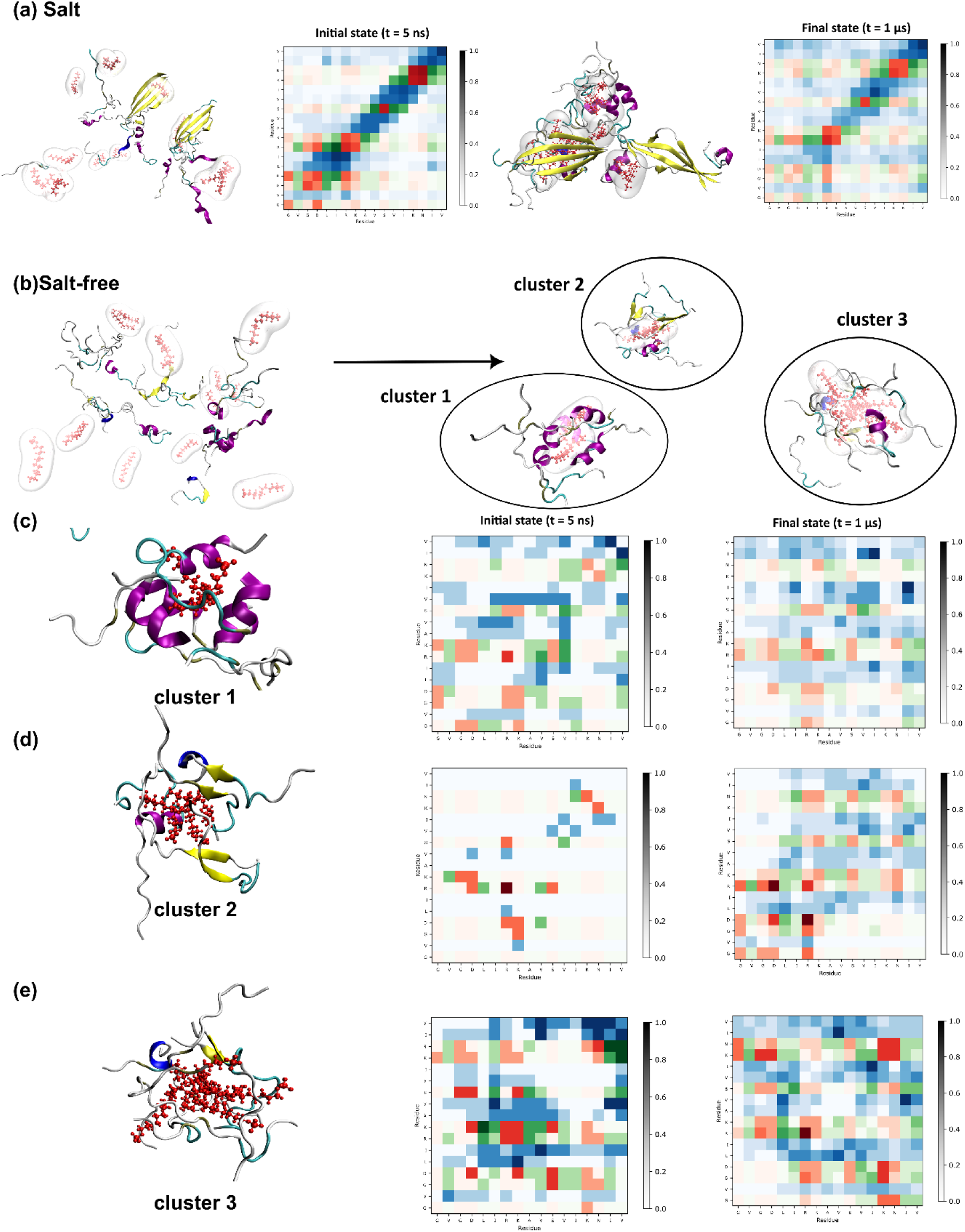
Inter-peptide contact maps and corresponding initial and final snapshots of peptide systems simulated under salt-containing and salt-free conditions, illustrating differences in aggregation behaviour and secondary structure evolution.(a) In the presence of salt, peptides form distinct aggregates over time. Initial and final snapshots are shown alongside residue–residue contact maps, highlighting the emergence of stable interactions associated with aggregation. (b) In the salt-free system, partial aggregation into small peptide clusters is observed at the end of the simulation. (c) Cluster 1, primarily composed of α-helical peptides, shows sparse initial contacts that become denser over time. As the helices form and SDS monomers insert between peptides, residue–residue contacts distribute along the full peptide length. However, the decreased color intensity in the contact map suggests these interactions are weaker and more transient. (d) Cluster 2, also from the salt-free system, exhibits contact patterns similar to the salt-containing system but aggregates more slowly. Interactions are predominantly localized near the N-terminus, likely indicating the formation of β-sheet-rich structures. (e) Cluster 3 consists of peptides with mixed secondary structure elements—partial helices, β-sheets, and predominantly random coils. Initial contacts are minimal, but strong aggregation is evident by the end of the trajectory. The contact map shows extensive interactions across the peptide length, though disordered in nature. Among the three clusters, the disordered cluster shows the highest number of contacts, followed by the β-sheet cluster (mainly N-terminal interactions), and then the helical cluster with the fewest contacts—likely due to the spatial separation introduced by SDS and the compact nature of helices reducing surface contact area. Hydrophilic and hydrophobic residues are represented in red and blue, respectively. Residue–residue contact maps use a color-coded scheme: red (hydrophilic–hydrophilic), blue (hydrophobic–hydrophobic), and green (hydrophilic–hydrophobic), with intensity indicating contact frequency.

**Figure S6.**
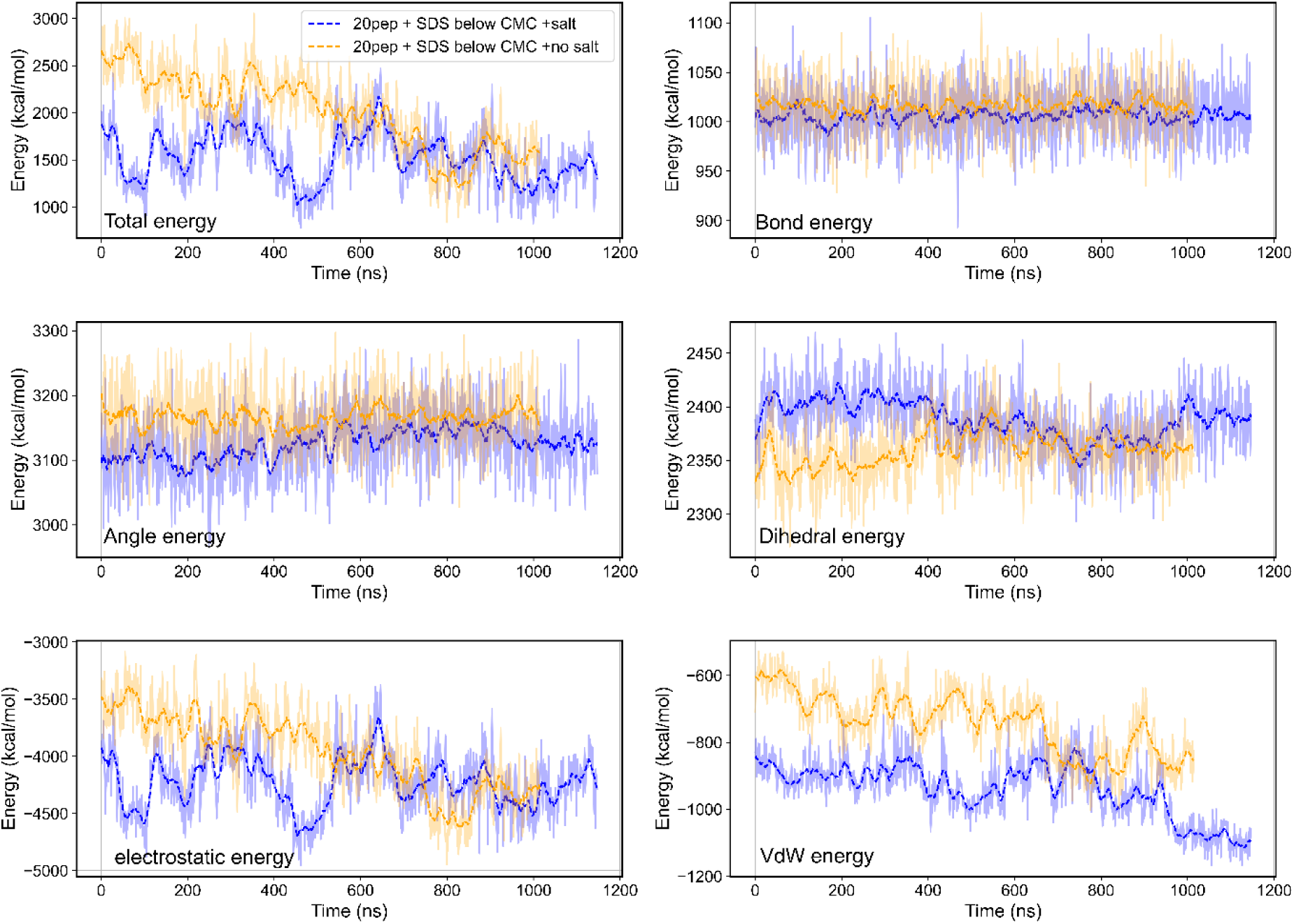
Comparison of different energy components over the simulation time for two 20-peptide systems: one with SDS monomers below CMC in the presence of salt (orange), and the other with SDS below CMC in the absence of salt (blue).Panels show the evolution of total energy, bond energy, angle energy, dihedral energy, electrostatic energy and van der Waals (VDW) energy.The salt-free system exhibits stronger electrostatic and van der Waals interactions, leading to a more negative total energy and larger energy fluctuations, consistent with enhanced peptide aggregation and clustering.In contrast, the salt-containing system shows reduced electrostatic interactions, resulting in more stable energy profiles and less compact aggregation. Bond and angle energies remain relatively stable across both systems, whereas dihedral energy suggests greater conformational constraint in the salt-free system due to tighter peptide packing.

**Figure S7.**
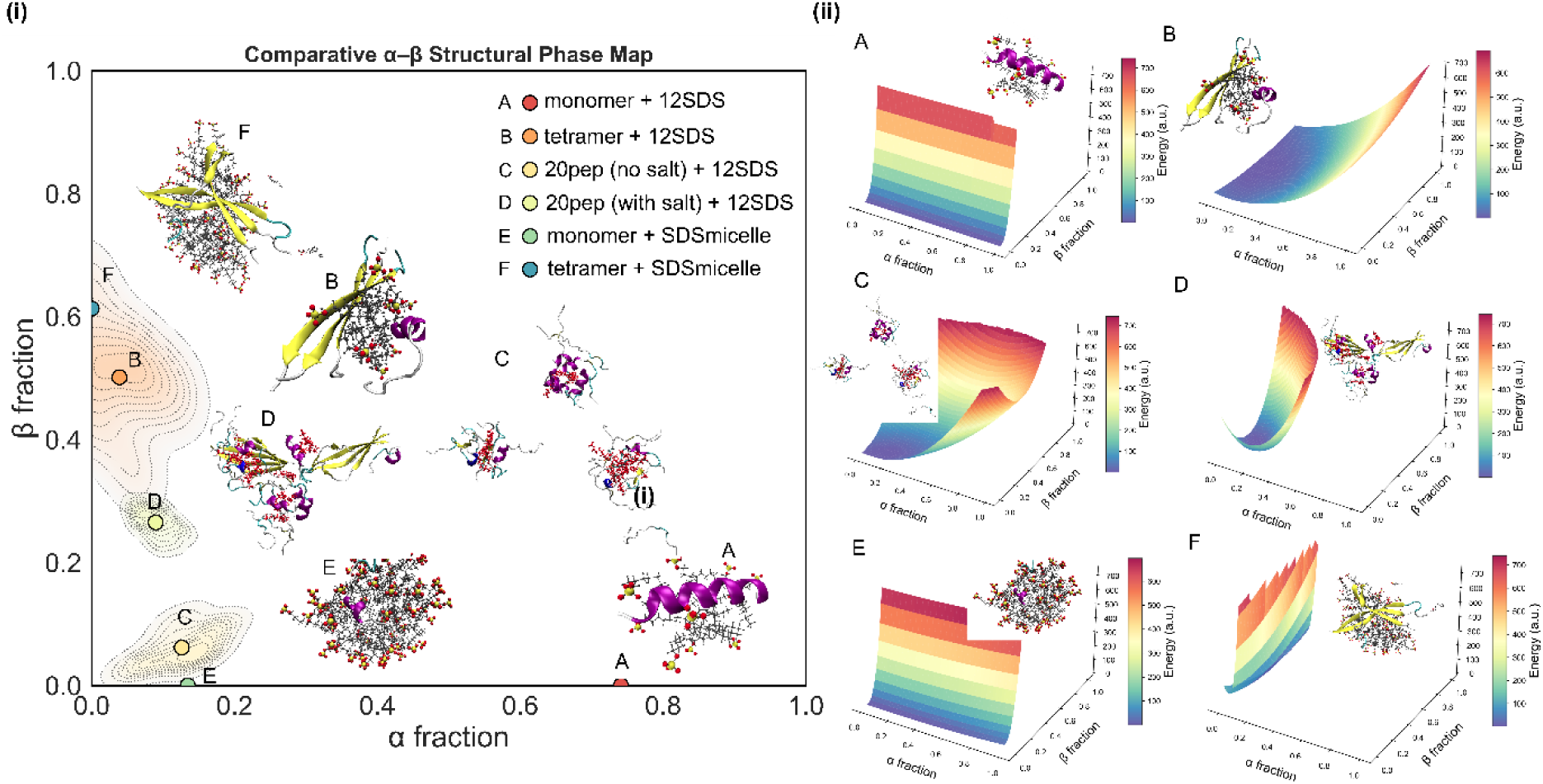
(a)Two-dimensional structural phase map showing the fractional α-helical (α fraction) and β-sheet (β fraction) content adopted by Uperin 3.5 under six distinct surfactant environments. Systems include: A. monomer + sub-CMC SDS, B. tetramer + sub-CMC SDS, C 20-peptide system without salt + sub-CMC SDS, D. 20-peptide system with salt + sub-CMC SDS, E. monomer + SDS micelle (above-CMC) and F. tetramer + SDS micelle. Representative structural snapshots are overlaid at their corresponding phase-space locations. (b)Right-hand panels show the underlying free-energy surfaces sampled by each system, illustrating how surfactant charge, concentration, peptide stoichiometry, and ionic strength reshape the accessible α/β conformational landscape. Together, these maps capture environment-driven structural polarization between α-helical antimicrobial peptide-like states and β-sheet-rich amyloidogenic states.

**Figure S8.**
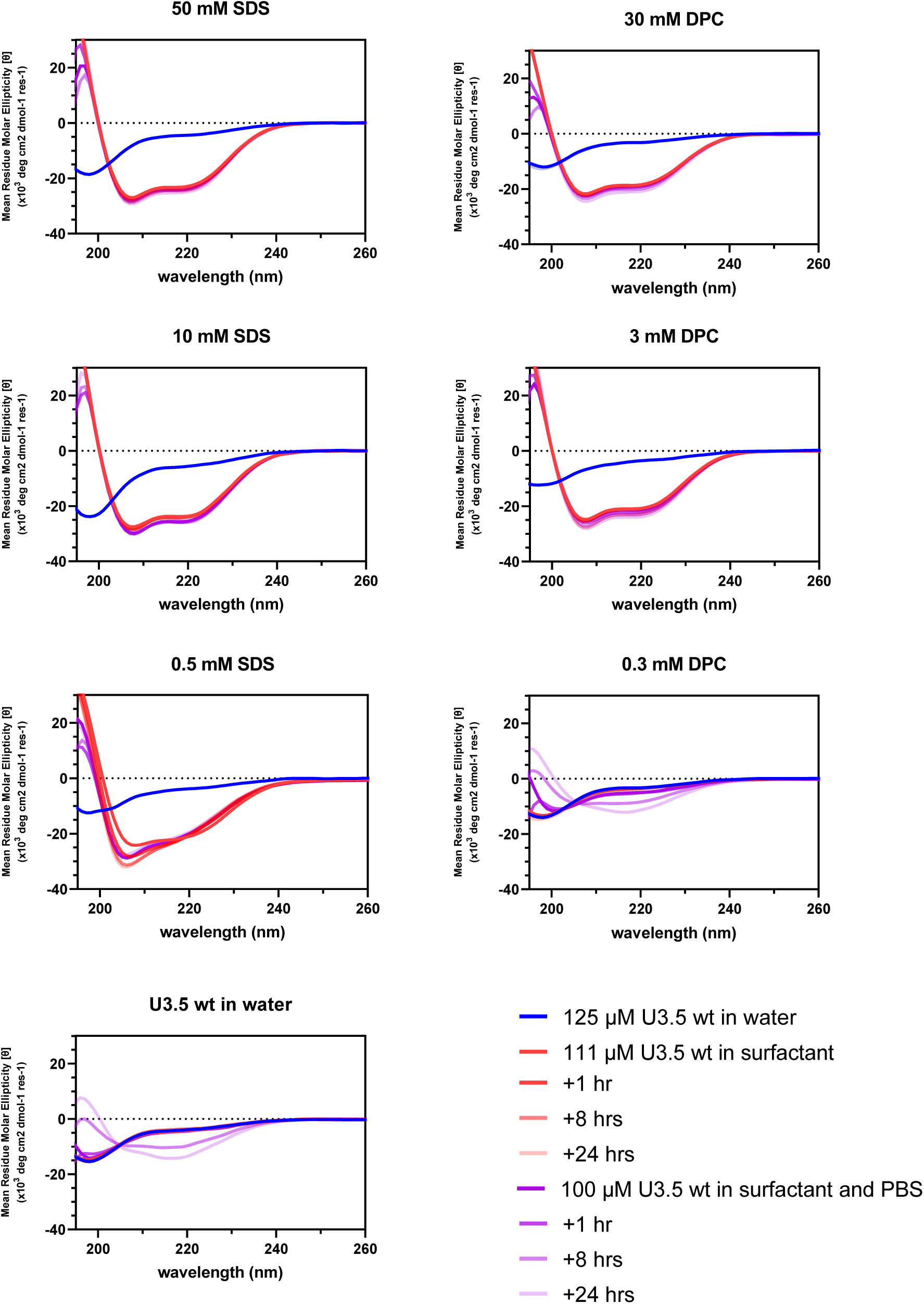
CD spectra of U3.5 self-assembly in the presence of surfactant. Measurements were conducted at 37 °C. Firstly, the peptides were measured in water for 1 hour to ensure no self-assembly is occurring. Then, each surfactant was added to their respective concentrations, and measured for 24 hours. Following this, 10x PBS was added as a 1 in 10 dilution, and the CD of the peptide ssamples were measured for further 24 hours.

**Figure S9:**
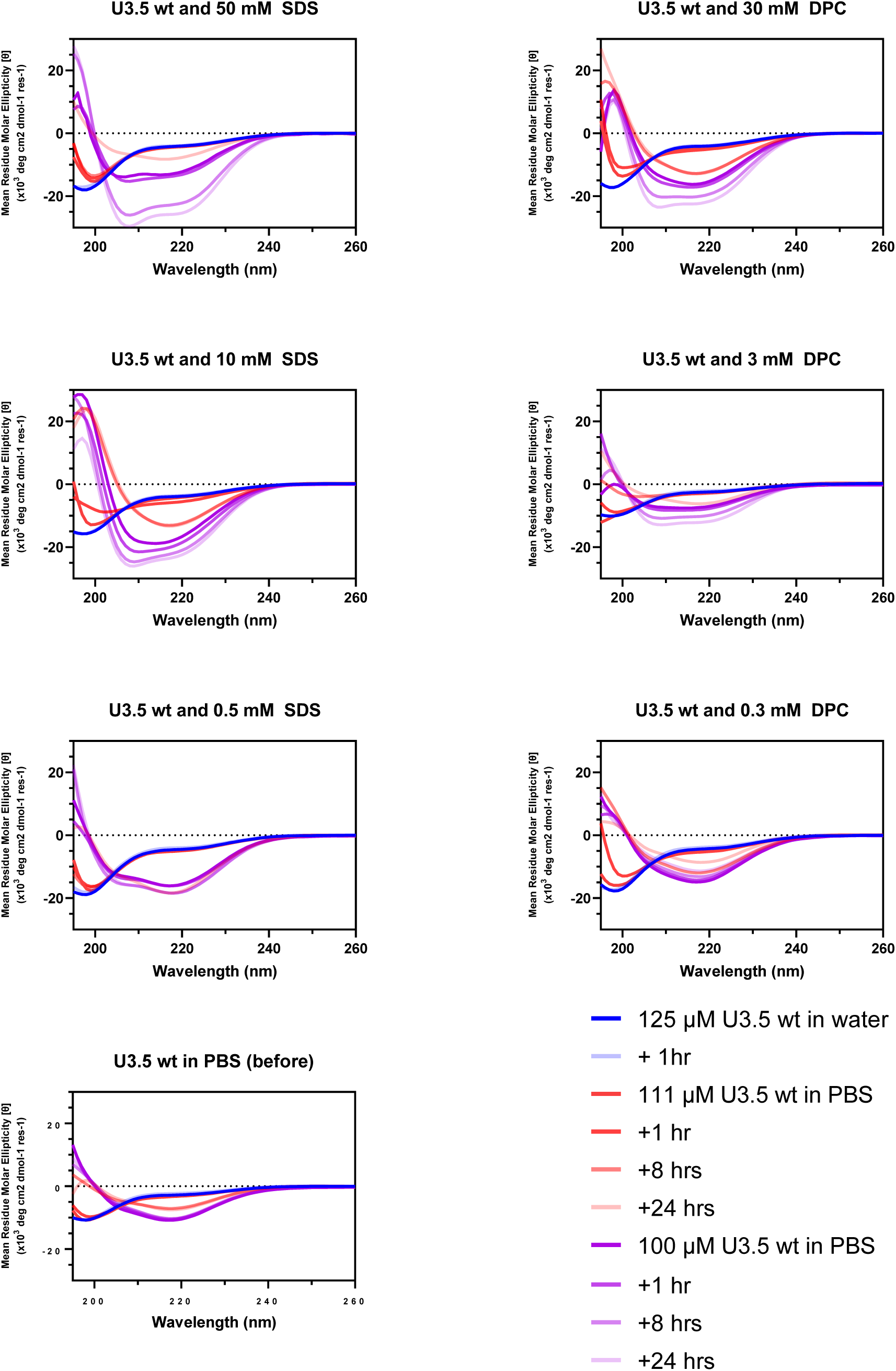
CD spectra of U3.5 wt disaggregation in the presence of surfactant. Measurements were conducted at 37 °C. Firstly, the peptides were measured in water for 1 hour to ensure no self-assembly is occurring. Then, 10x PBS was added as a 1 in 10 dilution, and the peptides were measured for another 24 hours. Following this, each surfactant was added to their respective concentrations, and measured for 24 hours.

